# Deep Learning Approaches to the Phylogenetic Placement of Extinct Pollen Morphotypes

**DOI:** 10.1101/2023.07.09.545296

**Authors:** Marc-Élie Adaïmé, Shu Kong, Surangi W. Punyasena

**Affiliations:** Department of Plant Biology, University of Illinois Urbana–Champaign, Urbana, IL 61801; Department of Computer Science and Engineering, Texas A&M University, College Station, TX 77843

## Abstract

The phylogenetic interpretation of pollen morphology is limited by our inability to recognize the evolutionary history embedded in pollen features. Deep learning offers tools for connecting morphology to phylogeny. Using neural networks, we developed an explicitly phylogenetic toolkit for analyzing the overall shape, internal structure, and texture of a pollen grain. Our analysis pipeline determines whether testing specimens are from unknown species based on uncertainty estimates. Features of novel specimens are passed to a multi-layer perceptron network trained to transform these features into predicted phylogenetic distances from known taxa. We used these predicted distances to place specimens in a phylogeny using Bayesian inference. We trained and evaluated our models using optical superresolution micrographs of 30 *Podocarpus* species. We then used trained models to place nine fossil *Podocarpidites* specimens within the phylogeny. In doing so, we demonstrate that the phylogenetic history encoded in pollen morphology can be recognized by neural networks and that deep-learned features can be used in phylogenetic placement. Our approach makes extinction and speciation events that would otherwise be masked by the limited taxonomic resolution of the fossil pollen record visible to palynological analysis.

**Significance Statement:** Machine learned features from deep neural networks can do more than categorize and classify biological images. We demonstrate that these features can also be used to quantify morphological differences among pollen taxa, discover novel morphotypes, and place fossil specimens on a phylogeny using Bayesian inference. Deep learning can be used to characterize and identify and morphological features with evolutionary significance. These features can then be used to infer phylogenetic distance. This approach fundamentally changes how fossil pollen morphology can be interpreted, allowing greater evolutionary inference of fossil pollen specimens. The analysis framework, however, is not specific to pollen and can be generalized to other taxa and other biological images.

## 1 Introduction

Machine learning in the computational biological literature has largely focused on biological classification and categorization, that is, developing neural networks for *K*-way classification, where *K* represents the number of taxa in a training set. Neural networks are used to identify features that define individual classes (taxa) and not the features that define the evolutionary relationships among taxa. This poses a challenge when classifying fossil specimens, many of which may be derived from extinct species that are entirely new to a trained network.

New methods are needed to detect novel fossil taxa and to accurately place them within a phylogeny. Deep learning techniques, such as convolutional neural networks (CNNs) are capable of extracting key discriminative features from a given image [23, 22]. Novelty can be identified by measuring the uncertainty within the *K*-way classification for a given specimen: high uncertainty implies a high likelihood that the specimen is from an unknown taxon [13, 20]. For phylogenetic placement, however, neural networks must explicitly incorporate evolutionary distances. Models must be trained not to identify the traits which define classes, but to recognize phylogenetic synapomorphies - derived features that are shared between sister taxa.

Our insight is that phylogenetic distances can be used to re-learn and re-weight morphological features derived from CNNs. CNN features can be treated as a vector of continuous morphological traits for placing taxa within a known phylogeny using Bayesian inference. We obtained these weighted features by first extracting the features from classification CNNs and then passing them to a second embedding model trained to transform features according to a phylogenetically-informed distance function. This is a fundamentally new approach in applying machine learning to biological classification.

Fossil pollen is an example of a fossil record where many morphotypes have unknown or uncertain biological affinities [26, 2, 39]. So although fossil pollen comprises one of the most extensive paleontological archives of terrestrial ecosystems, poor taxonomic resolution has limited evolutionary and paleoecological inference [26, 33]. Our approach builds on our previous work demonstrating that well-constructed CNN analyses can transfer learning from modern pollen morphology to fossil [36] and expands the range of questions that can be addressed by pollen data, offering new insights into paleoecology, plant evolution, and phylogenetics. It allows us to recognize extinct morphotypes and reconstruct their phylogenetic relationship to extant taxa. It allows us to reassess and recalibrate existing phylogenetic trees using microfossil data. The development of phylogenetically informed deep learning models is not specific to pollen, but also can be generalized to other taxa with robust phylogenies and standardized morphological imaging.

### 1.1 General analysis pipeline

#### 1.1.1 Image classification

We began by training three independent convolutional neural networks (CNNs) for *K*-way classification by exploiting three individual modalities (shape, internal structure, and texture) (Fig. 1A). The first, holistic CNN (H-CNN) took as input maximum intensity projections (MIPs) of the whole grain. The second, cross-sectional CNN (C-CNN) used axial slices taken through the entirety of a pollen grain. The third, patch CNN (P-CNN), was trained on overlapping square crops of the initial MIP image, each covering ∼10% of the entire grain. The result of each CNN was a *K*-dim vector of classification scores that captured probabilities for each known taxonomic classification. We used a probabilistic fusion model (FM) (Fig. 1B) to combine these single-modal probabilities into a fused multimodal probability for better performance.

**Figure 1.**
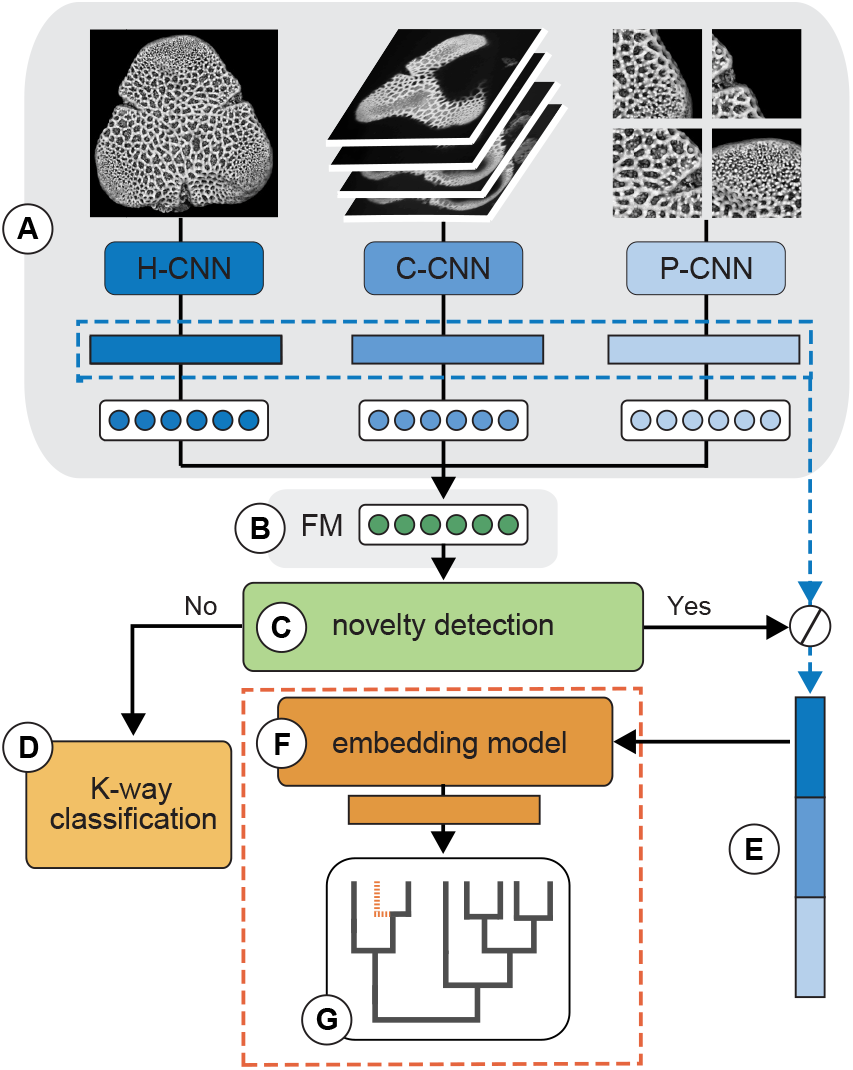
Flowchart illustrating the trained multimodal neural network pipeline. Three representations of superresolution images are passed through CNNs capturing shape, internal structure, and texture (H-CNN, C-CNN, P-CNN; A). The three sets of classification scores are fused (fused model, FM; B) and the analysis determines whether a specimen belongs to a known taxon by assessing the network’s uncertainty during image classification (ROC analysis; C). If the specimen is recognized as one of the *K* taxa with high confidence (low uncertainty), its predicted taxon is reported (D). Otherwise, its features are extracted across all three image modalities and concatenated (E) as input to a multi-layer perceptron (F), which is trained to transform these features to an embedding feature to better compute phylogenetic distances from known taxa. Embedding features are clustered with the features of known taxa for phylogenetic placement, based on Bayesian inference (G).

#### 1.1.2 Novel taxon detection

The total uncertainty of each CNN and FM classification was calculated using entropy computed over the product of their classification probabilities. We used receiver-operating characteristic (ROC) analysis [42] to establish the uncertainty threshold above which a specimen should be considered novel (Fig. 1C). ROC is widely used to evaluate the performance of binary classification tasks [42, 16]. The area under the ROC curve (AUROC) [20] quantified our models’ ability to discriminate between known and novel pollen taxa. We determined the threshold that optimized both the precision and recall of novel taxa. If the model’s uncertainty score was below this threshold, the specimen’s most likely predicted taxon was reported (Fig. 1D). Otherwise, the sample was flagged as novel, and the features extracted from the three classification CNNs (Fig. 1E) were forward passed through an embedding model for phylogenetic placement (Fig. 1F).

#### 1.1.3 Phylogenetic placement

Our phylogenetic embedding model is a trained multi-layer perceptron (MLP) that transforms a specimen’s classification features to embedding features based on a learned phylogenetic distance function (Fig. 1F). The resulting embedding features capture the phylogenetic information within classification features. The embedding features can then be used to place a novel specimen on an established phylogeny (Fig. 1G). We train the MLP over specimens from *K* known taxa by minimizing the differences in the computed Euclidean distances between pairs of specimens’ embedding features and ground-truth phylogenetic distances. We then use Bayesian inference to reconstruct the original tree topology from the continuous machine-learned features, applying the Brownian motion (BM) model for character evolution [8, 9, 30].

## 2 Materials and Methods

### 2.1 Specimens and molecular phylogenies

We used 30 extant *Podocarpus* (Podocarpaceae) species and nine fossil *Podocarpidites* specimens (Fig. 2) [34] and a fossil-calibrated phylogeny [24] to train and evaluate our neural networks. Time-calibrated phylogenies allowed us to work with absolute ages (in millions of years, Ma) rather than relative phylogenetic distances.

**Figure 2.**
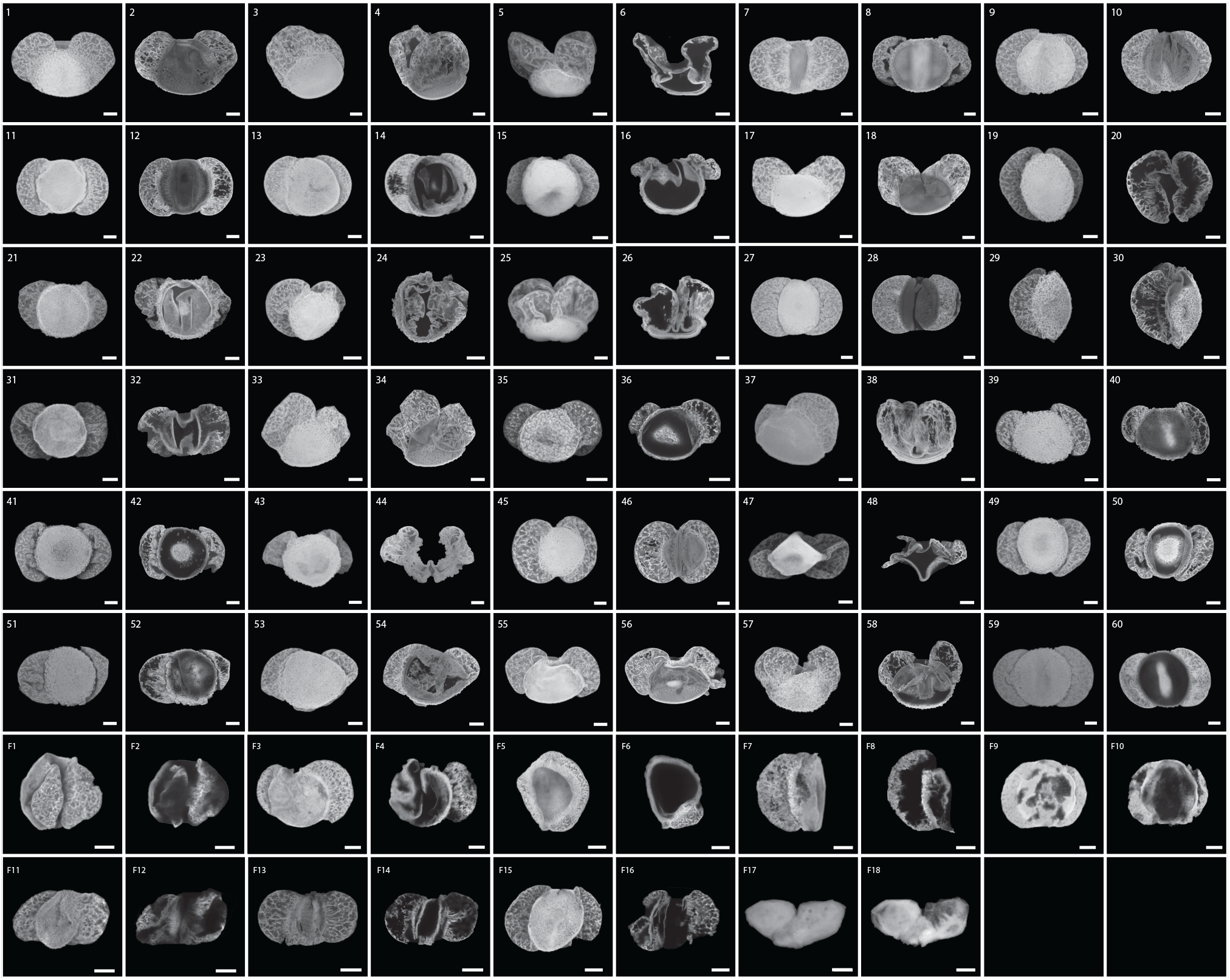
Modern and fossil specimens of *Podocarpus*. Each specimen is represented by a pair of images consisting of a maximum intensity projection (MIP) (first image) and a cross-sectional image (second image). Plate numbers and their corresponding species can be found in Supplemental Tables S4 and S5 (scale bar: 10 µm).

Specimens were from the collections of Smithsonian Tropical Research Institute and Utrecht University. The pollen image dataset included 309 specimens from Australasia/Indomalaya (16 species), the Neotropics (10 species), Africotropics (2 species), and Asian Palearctic (1 species) (Supplemental Table S4). The dataset also includes nine fossil *Podocarpidites* specimens from the Late Cretaceous and Neogene of Colombia, Peru, and Panama (Supplemental Table S5) [17, 27, 4]. Modern taxon names were reviewed and amended according to the taxonomic nomenclature in Tropicos [41]. Specimens were imaged with Airyscan confocal superresolution microscopy at 630x magnification (63x/1.4NA oil objective, 0.08 µm per pixel resolution) as a series of axial focal planes (Z-stacks) at 0.19 µm increments [34]. We manually cropped and masked grains to remove background debris.

For the phylogenetic model, we relied on a conifer-wide fossil-calibrated phylogeny based on a concatenated dataset of two chloroplast DNA (cpDNA) protein-coding genes, rbcL and matK, and the 18S nuclear ribosomal DNA (rDNA) gene [24]. The reconstruction used a partitioned maximum likelihood analysis and assumed a GTR+Γ nucleotide substitution model. Most clades were well-supported, with high bootstrap support. Of the 45 species in the *Podocarpus* image dataset [34], only 30 were included in the phylogeny and were therefore selected for this study.

### 2.2 Image modalities

For our first imaging modality, we generated images of the external structure and shape of an entire pollen grain using maximum intensity projections (MIP) of the entire specimen image stack [33]. We applied contrast-limited adaptive histogram equalization (CLAHE) to standardize all images [46].

For the second modality, we used cross-sectional images. For the modern specimens, we generated a series of internal cross-sections by producing MIPs of staggered subsections of the image stack [36]. Each substack included 15 axial cross-sectional planes (2.85 µm depth), offset by 10 planes (1.90 µm). Because our fossil specimens were highly compressed, instead of subsections we used each individual plane of the image stack as our cross-sections.

The final modality was image patches that sampled a small portion of the grain. The whole-grain MIP for each specimen was divided into 10-13 smaller square patches, each of which covered *∼*10% of the original MIP image and overlapped *∼*25% with adjacent patches [19].

### 2.3 Deep learning models

#### 2.3.1 Network architectures

We trained three separate *K*-way classification CNNs for each of our three modalities: a holistic image CNN (H-CNN), a cross-sectional CNN (C-CNN), and a patch CNN (P-CNN). All three CNNs incorporated ResNeXt-101 architecture [44] with the exception of the last layer, which was modified to output *K*-dim logit vectors. We normalized the logit vectors using softmax to produce *K*-dim probability vectors, which indicated the probability of the input image being classified as each of the *K* pollen taxon classes.

We fine-tuned the ResNeXt CNNs (pre-trained on ImageNet images) using our modern reference pollen dataset [7, 11]. For a given specimen, the H-CNN produced a *K*-dim logit vector (i.e., the feature prior to the *K*-dim softmax scores). The C-CNN and P-CNN used multiple cross-sections or patches and produced multiple logit vectors, respectively. We next computed the mean of the per-patch logit vectors and the mean of the per-cross-sectional logit vectors. Lastly, we summed the three logit vectors and normalized by softmax to obtain a *K*-dim classification probability vector.

#### 2.3.2 Training details

Specimens of the *K* known taxa were randomly split into training (70%) and validation (30%) sets. We augmented the training data by adopting common augmentation techniques including random flip, random rotation (with probability *p*=0.5 by degrees in [-90^*◦*^, +90^*◦*^]), and random translation (within pixel displacement in [-30, +30]). We resized the augmented images to 224 × 224 × 3 pixel resolution using bilinear interpolation.

We trained the *K*-way classification CNNs by minimizing the cross-entropy loss:

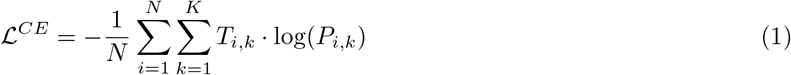

where *N* is the number of specimens in the training set, *K* is the number of taxa, and *P*_*i,k*_ is the softmax probability of specimen-*i* classified as taxon-*k. T*_*i,k*_ is a binary indicator:

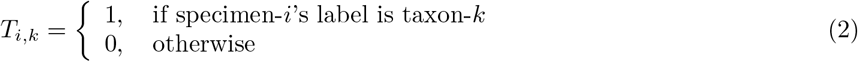

We used the stochastic gradient descent optimizer with momentum (0.9). We adopted a stage-wise learning rate schedule with an initial learning rate of 0.0009, decreased by half every two epochs. All models were trained for 20 epochs with a batch size of 10.

#### 2.3.3 Fused model (FM)

Given a specimen *X* and its three modalities (*x*_1_, *x*_2_, and *x*_3_), we fused the predictions of the three modalities by assuming conditional independence given the class label [5], i.e., *p*(*x*_1_, *x*_2_, *x*_3_|*y* = *k*) = *p*(*x*_1_|*y* = *k*) *· p*(*x*_2_|*y* = *k*) *· p*(*x*_3_|*y* = *k*). Further, we assumed a uniform prior, i.e., *p*(*y* = 1) = *…* = *p*(*y* = *K*). Below is the fusion result:

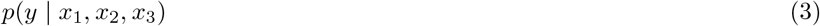

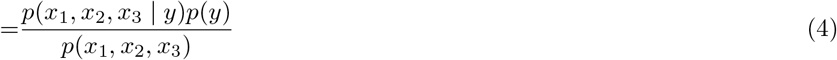

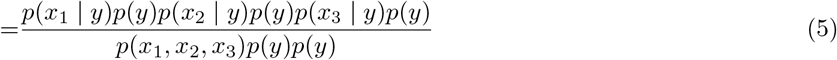

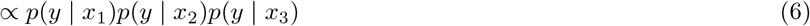

(4) applies the Bayes’ theorem, (5) assumes conditional independence, (6) assumes uniform prior w.r.t class labels. To derive the probability after fusion, we normalized (6) to sum-to-one over the *K* classes, so that:

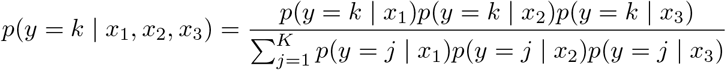

#### 2.3.4 Model validation

We assessed our models using five random splits of training (70%) and validation (30%) image sets. We report the mean of per-taxon accuracies. Confusion matrices were presented to provide a detailed report of per-taxon accuracies (Fig. 3).

**Figure 3.**
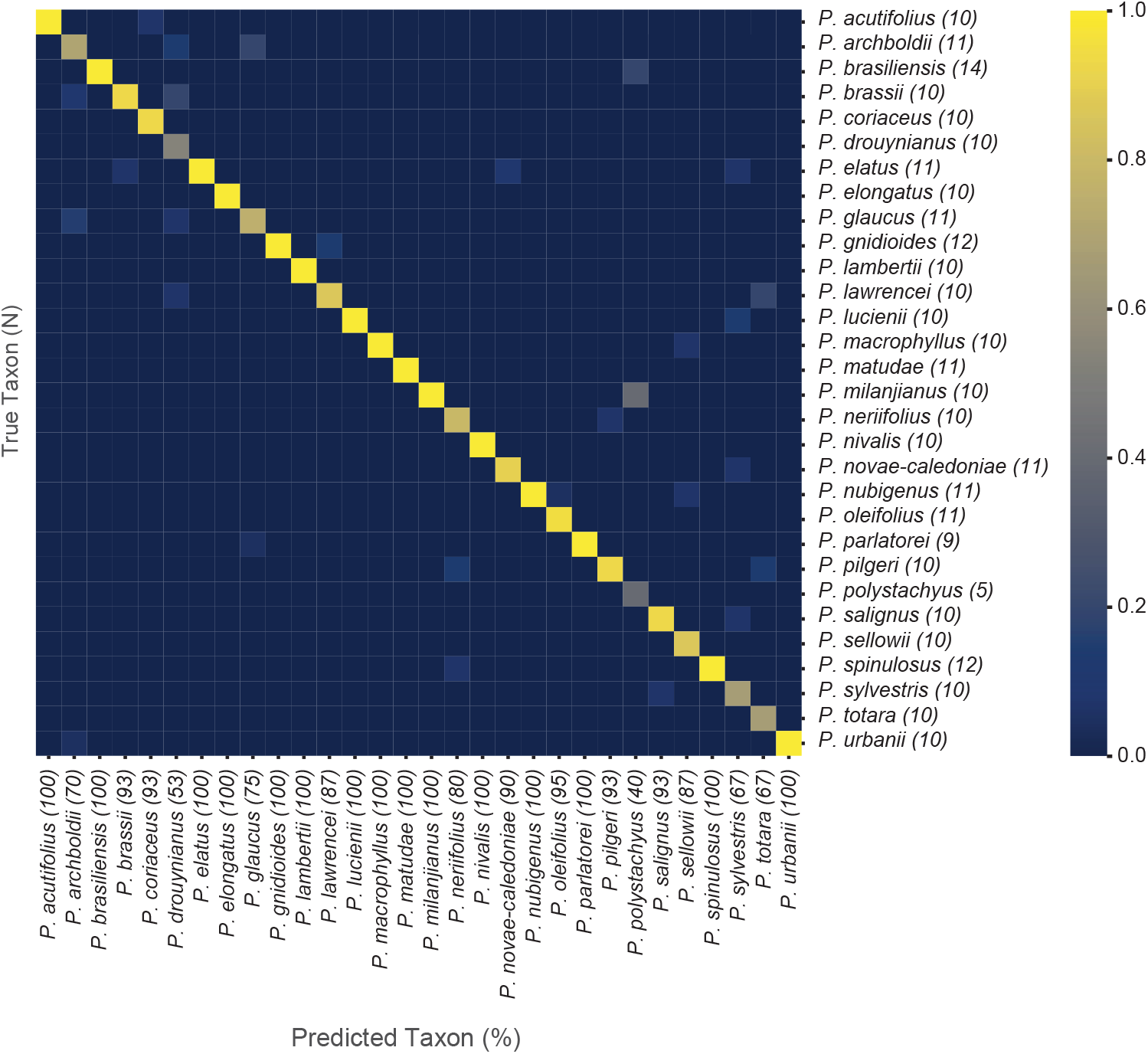
Average confusion matrix depicting the classification accuracy of the fused model (FM) for the modern *Podocarpus* species. Rows correspond to the true taxon while columns represent the model’s predictions.

### 2.4 Novelty detection

#### 2.4.1 Uncertainty estimates

We used entropy [16] to assess whether a specimen should be considered as novel to the trained model. Given specimen *X* and its probability of being classified as taxon-*k, p*(*y* = *k* | *X*) for *k* = 1, …, *K*, we computed entropy *H*(*p*) as:

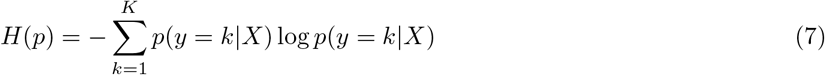

We compared novelty detection performance in each of the three CNN models and the FM. To measure the models’ abilities to determine true novel morphologies, we used the Receiver Operating Characteristic (ROC) curve between True Positive Rate (TPR) and False Positive Rate (FPR). ROC is a graph showing the performance of a binary classification model at different thresholds. We computed the area under the ROC curve (AUROC) as the summary number to benchmark methods. Higher AUROC means better performance for novelty detection [20].

#### 2.4.2 Optimal threshold selection

Although AUROC effectively summarizes model performance with varied thresholds, real-world systems require a single operation threshold. We used the method introduced in [42] to find an optimal threshold *c*^*∗*^ for fossil pollen analysis. We analyzed the ROC curve to find *c*^*∗*^, using the TPR and FPR for a binary classification model designed to detect specimens of novel pollen types. We evaluated the performance of the model by using our training set to represent the known specimens and the pseudo-novel set to represent the novel specimens. We optimized the following objective function [42] to obtain *c*^*∗*^, the value at which TPR and (1-FPR) values are closest to AUROC while minimizing their absolute difference:

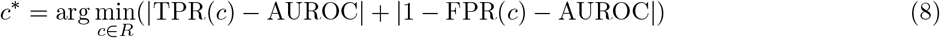

### 2.5 Phylogenetic placement

#### 2.5.1 Model and loss function

Specimens identified as novel were placed within an existing phylogeny using a trained embedding model, implemented as a multi-layer perceptron (MLP). MLPs are a special form of CNNs in which all the convolution layers have 1 ×1 kernels. For a novel specimen, the input to the MLP was a concatenation of the specimen’s three classification feature vectors (with 2048 dimensions each), extracted from the three classification CNNs. The MLP outputs a low-dimensional embedding feature vector **v ∈** *R*^256^ used to place the specimen within a phylogeny.

Our MLPs have five convolutional layers. The first layer learns three non-negative parameters to weigh the three classification features learned by individual CNNs. Between every two consecutive layers, we inserted a rectified linear activation unit (ReLU) for nonlinearity [29] and dropout layers (with a dropout rate of 0.2 to prevent overfitting [38]. The complete network architecture is described in the supplementary materials (Supplemental Fig. S5 and Table S6).

Our MLP is lightweight relative to a CNN, allowing us to train with all training specimens’ features as a single batch. We used the Adam optimizer [18] with a constant learning rate of 0.00001 and a weight decay of 10^*−*4^. We trained the MLP for 20,000 iterations (∼10 min running time on a GPU).

We developed a simple loss function to train the MLP to learn phylogenetic patterns from classification features. The loss function minimizes the error between ground-truth and predicted evolutionary distances separating pairs of specimens. Because the MLP outputs an embedding feature *v*_*i*_ for specimen-*i*, between specimen *i* and *j*, we compute their Euclidean distance.

With the known ground-truth phylogenetic distance *D*_*i,j*_, we minimize the mean square error (MSE) between *D*_*i,j*_ and 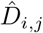 :

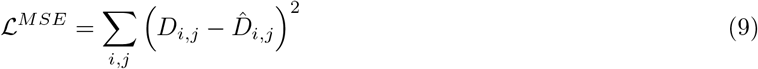

All the ground-truth and predicted phylogenetic distances in *D* and 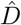 were re-scaled to the range [0, 1].

#### 2.5.2 Tree construction

We constructed phylogenetic trees from the transformed features using Bayesian inference and assuming the Brownian motion (BM) model for character evolution [8, 9, 30]. Analyses were performed using RevBayes [14]. We performed Markov Chain Monte Carlo (MCMC) analyses consisting of two independent runs with four chains each, running for 1 million generations. Trees were sampled every 1000 generations, and initial pre-stationarity generations were discarded according to the burn-in value determined with Tracer (v.1.7.1). Tree files from both independent runs were combined using LogCombiner (v.1.10.4) to generate the final phylogeny. The maximum clade credibility (MCC) tree, which maximizes the posterior probability of each individual clade, was chosen as the final model using TreeAnnotator (v.2.4.2). The resulting trees were visualized and edited using the functions plotTree and plotFBDTree in the R package “RevGadgets” (v.1.1.1) [40]. Branches in the tree with a posterior probability *≥* 0.95 were considered significantly supported.

### 2.6 Pseudo-Novel evaluation experiments

#### 2.6.1 Evaluation of novelty detection

To evaluate our methods for discriminating among known and novel taxa, we adopted a “leave-one-out” protocol [21], where a taxon was excluded from the training dataset and its specimens were treated as novel morphotypes. We ran five separate experiments and selected one of five taxa from biologically meaningful subclades in the reference phylogeny: *P. drouynianus, P. elongatus, P. neriifolius, P. oleifolius*, and *P. totara*.

We forward-passed all specimens of a pseudo-novel taxon and the known validation specimens to their corresponding trained CNNs. We computed entropy over their class probability distributions and ran a ROC analysis. The analysis provided an assessment of the model’s performance in novelty detection. In each run of these experiments, we also computed an optimal threshold. We averaged these thresholds to obtain a single threshold for identifying new fossil morphotypes.

#### 2.6.2 Evaluation of phylogenetic placement

We extracted the CNN-learned features of all pseudo-novel specimens and passed them to the trained MLPs. We then concatenated the resulting output features with the average feature results for the known species. This produced a continuous character matrix of pseudo-novel specimens and known species. We used the character matrix to construct Bayesian inference trees under a single rate Brownian motion model. The trees were simulated under a birth-death model, with all parameters defined following [30]. Because our modern dataset sampled 30% of extant *Podocarpus* taxa, the sampling rate was set to 0.3.

### 2.7 Fossil application

We forward-passed the nine fossil *Podocarpidites* specimens through our fully trained CNN-MLP pipeline, where the model had been trained on all modern specimens. This produced a continuous character matrix of fossil specimens and extant species, which were then used to construct a Bayesian inference tree. We performed tip-dating under the fossilized birth-death (FBD) process, enabling the simultaneous analysis of both modern and fossil morphotypes while calibrating divergence times estimates in a Bayesian framework [12, 45]. Tip ages were included as single-point occurrence times corresponding to the most likely ages of the fossil specimens. The diversification and turnover rates were drawn from a uniform distribution over an interval ranging from 0 to 1. Speciation was defined as the sum of diversification and turnover rates, while extinction was set to be equal to the turnover rate. The root age was drawn from a uniform distribution with a lower bound of 67.2 Ma, corresponding to the age of our oldest fossil, and an upper bound of 82 Ma, based on the maximum root age estimated in [35]. Similarly, the fossil rate was drawn from a uniform distribution ranging from 0 to 1. The sampling fraction of extant taxa was set at 0.3. The resulting time-calibrated MCC tree provided us with the most likely placements of our fossil specimens within the reference phylogeny, accompanied by their corresponding posterior probabilities.

## 3 Results

### 3.1 Baseline classification accuracy

We measured taxon classification accuracies for the three imaging modalities (H-CNN, C-CNN, and P-CNN), and the fused model (FM). The FM and C-CNN produced the highest average accuracies, 90.40% and 90.60% average accuracy, respectively. The P-CNN scored lower (79.80%) and the H-CNN lower still (55.00%) (Fig. 3, Supplemental Fig. S1, Supplemental Table S1).

### 3.2 Novel taxon detection

The C-CNN (*µ*=0.9425) and FM (*µ*=0.8987) markedly outperformed the P-CNN (*µ*=0.8407) and H-CNN (*µ*=0.7104) in detecting novel pollen morphotypes. The FM had the smallest variability overall (*σ*=0.0325). The P-CNN and H-CNN had higher variability (*σ*=0.0725 and 0.0591, respectively) (Supplemental Fig. S2 and Table S2).

### 3.3 Phylogenetic placement of pseudo-novel specimens

The Bayesian inference trees constructed from the learned MLP features largely replicated the *Podocarpus* molecular phylogeny and faithfully reconstructed two well-supported subgenera (*Podocarpus* subg. *Podocarpus* and *P*. subg. *Foliolatus*), with strong branch support for most clades across the trees (Supplemental Figs. S3.1-S3.5). The MLP determined the optimal weights for combining features from the three modalities (H-CNN, C-CNN, and P-CNN). In all five pseudo-novel experiments, the MLPs learned nearly equal weights for each modality (Supplemental Table S3).

For three pseudo-novel taxa *P. drouynianus, P. oleifolius*, and *P. totara*), all specimens were placed in their correct respective subclade, with high support values (P=1 in all three experiments) (Supplemental Figs. S3.1, S3.4, and S3.5).

For *P. elongatus*, the accuracy of the placement varied by specimen (Supplemental Fig. S3.2). Six specimens were correctly placed with *P. milanjianus*, with fairly high support (P=0.935). Three were placed elsewhere within subgenus *Podocarpus* and the remaining one was placed within subgenus *Foliolatus*. In the *P. neriifolius* experiments, seven specimens were correctly placed within the subclade comprising *P. brassii, P. archboldii, P. macrophyllus*, and *P. pilgeri*, with fairly high support (P=0.910) (Supplemental Fig. S3.3). The remaining three specimens were placed as sister to the entire *Foliolatus* subgenus.

### 3.4 Fossil application

All nine fossil *Podocarpidites* specimens were placed with *Podocarpus* subgenus *Podocarpus* (Fig. 4). Notably, the fossil placements were less certain than the pseudo-novel placements (Fig. 4 and Supplemental Figs. S3.1-S3.5).

**Figure 4.**
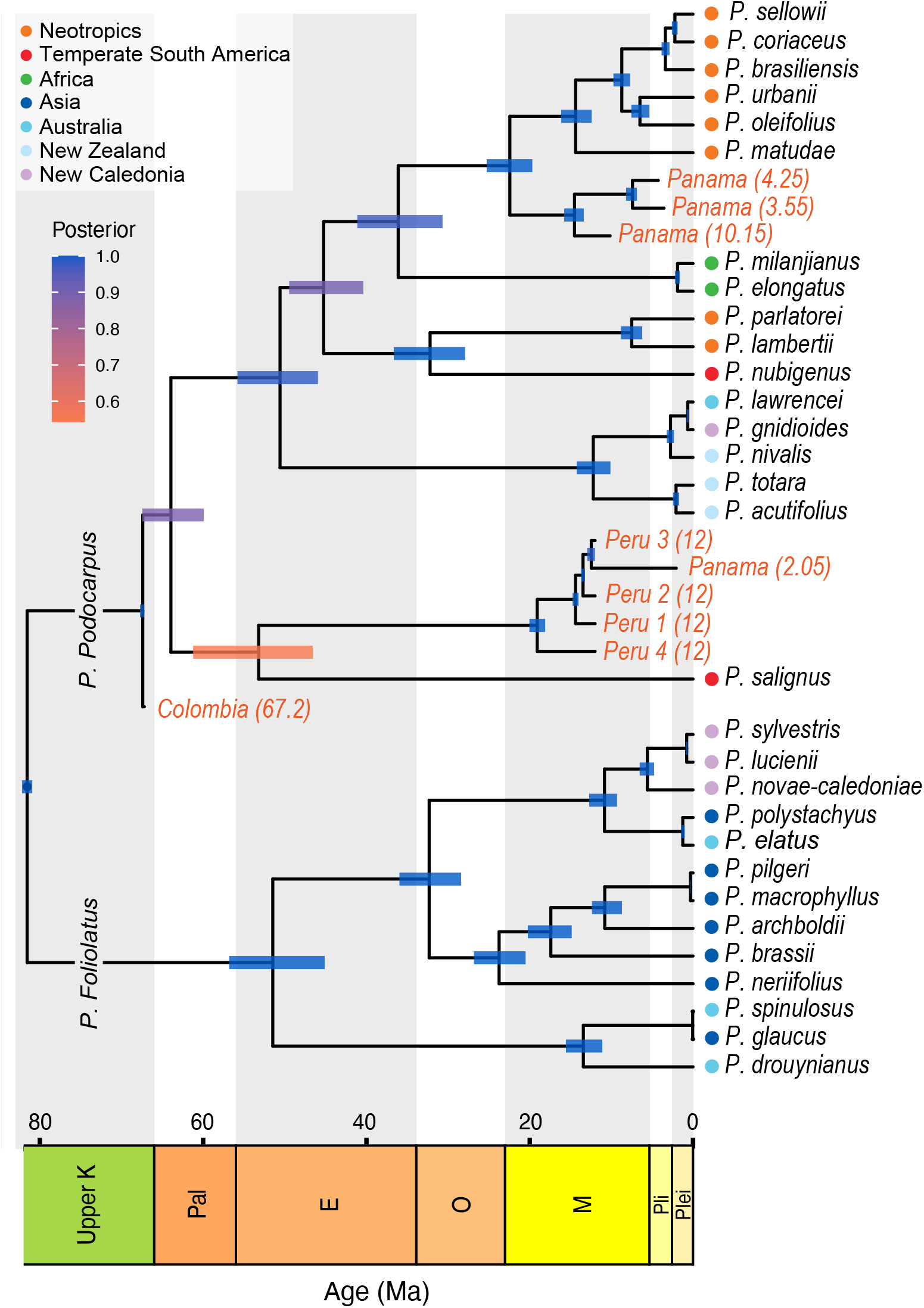
Combined tip-calibrated fossil tree simulated under the fossilized birth-death process (FBDP) using the MLP-transformed features. The tree topology replicates that of the reference phylogeny proposed by [24]. Branch lengths were estimated based on our fossil tip ages. The three oldest Panamanian specimens are placed as sister to the Central-South American tropical clade formed by *P. sellowii, P. coriaceus, P. brasiliensis, P. urbanii, P. oleifolius*, and *P. matudae*, while the youngest specimen is placed with the Chilean *P. salignus*. Likewise, the four Peruvian specimens are are placed with *P. salignus*. Finally, the late Cretaceous Colombian specimen is placed as sister to the subgenus *P. Podocarpus*.

The oldest specimen (Colombia, 67.2 Ma) was placed as sister to the entire subgenus, with strong support values (P=1). The three oldest Panamanian specimens (10.15, 4.25, and 3.55 Ma) were placed as sister to the Neotropical clade formed by *P. matudae, P. urbanii, P. oleifolius, P. brasiliensis, P. sellowii*, and *P. coriaceus* (P=1).

The youngest fossil and most taphonomically altered fossil (Panama, 2.05 Ma) was placed as sister to the Andean Chilean species *P. salignus*. The four Peruvian specimens (12 Ma) were also placed as sisters to *P. salignus*. The *P. salignus* and *Podocarpidites* clade was poorly supported (P=0.5426).

## 4 Discussion

CNNs are powerful tools for the analysis of biological morphology. Our results and previous research demonstrate their ability to capture the variability of pollen morphology and accurately classify species, even when there are limited morphological differences [33, 36, 3, 37]. However, our study takes the application of machine learning in biological classification further, demonstrating that features captured by neural networks can be interpreted and used within a phylogenetic framework. Our results show that machine-learned features can be used to interpret pollen morphology in an evolutionary context and place fossil specimens within a phylogeny.

Molecular phylogenetic studies place the origin of *Podocarpus* in the Late Cretaceous [35] or Paleocene [24, 1, 25], with more uncertain timing for the divergence of the two subgenera, *P. Podocarpus* and *P. Foliolatus*. The underlying uncertainty in published phylogenies introduced noise into our MLP models. Not all nodes of the original reference tree were well supported or fully resolved [24] and there were disagreements between the phylogeny used to train our models [24] and an alternative recent phylogeny [35]. Most notably, the African clade within *P. Podocarpus* is more closely related to the tropical Central-South American clade in our reference phylogeny [24], while in [35], the clade is more closely related to subtropical South American species. Additionally, in [35], the temperate South American species *P. nubigenus* and *P. salignus* are more closely related to the Australasian taxa of *P. Podocarpus* (*P. nubigenus, P. lawrencei, P. acutifolius, P. totara, P. gnidioides*, and *P. nivalis*). In contrast, in our reference phylogeny, *P. salignus* is placed as sister to the subgenus *P. Podocarpus* and *P. nubigenus* is more closely related to the tropical-subtropical South American species of *P. parlatorei* and *P. lambertii* [24].

*Podocarpus* pollen is bisaccate with large sacci that are coarsely endoreticulate. There is little morphological diversity within the genus, and palynologists generally do not attempt to identify individual species [31, 15, 28]. Despite this, we achieved high classification accuracies for our modern species with few per-species examples. Only the H-CNN performed poorly, with an average accuracy of 55.00% (Supplemental Table S1). In contrast, cross-sectional images and patches expanded the number of training examples per species and circumvented the “few-shot” learning problem of training models on a limited number of examples [6, 43].

Many misclassifications were of closely related species (Fig. 3, Supplemental Fig. S1). *P. archboldii, P. lawrencei, P. neriifolius, P. sylvestris, P. drouynianus, P. pilgeri*, and *P. totara* were confused with their sister species or species within the same subclade. Several misclassifications, however, were among distantly related species. *P. polystachyus* was misclassified as *P. milanjianus* and *P. brasiliensis*, likely due to its small sample size (n=5). Additionally, *P. glaucus* was confused for *P. archboldii* and *P. parlatorei, P. drouynianus* was confused for *P. brassii, P. archboldii*, and *P. lawrencei*, and *P. totara* was confused for *P. pilgeri*. Although we cannot identify specific morphological features which explain these misidentifications, we suspect that they are due to similarities in the endoreticulate patterning of their sacci, resulting from convergence or homoplasy.

We were able to successfully establish an uncertainty threshold for identifying taxa that were new or unknown to the model. The C-CNN and FM were the least variable in recognizing pseudo-novel specimens, with consistently high AUROC values. We expected the internal structure of the pollen wall to retain phylogenetically informative characters for identifying novel morphotypes based on prior research [36]. The P-CNN produced both the highest (i.e., *P. oleifolius*) and lowest (i.e., *P. elongatus*) AUROC values and was, therefore, the least reliable in detecting new species.

MLP-transformed CNN features closely reproduced the molecular phylogeny used to train our models. We correctly placed 43 out of 51 pseudo-novel specimens (Supplemental Figs. S3.1-S3.5). While we can not explain the inaccurate placement of the four incorrectly placed *P. elongatus* specimens, the sacci of the three erroneously placed *P. neriifolius* grains were obscured by their orientation. This may have led to their inaccurate placement in the reference phylogeny.

Our fully trained models also produced plausible placements of fossil specimens. The fossil pollen morphospecies *Podocarp-idites* is associated with *Podocarpus*, although other taxa - namely the extant New Zealand species *Lepidothamnus laxifolius* and Laurasian Paleogene Pinaceae species - have similar morphologies [28, 10]. Our *Podocarpidites* fossils are all Neotropical and Neogene or younger, with the exception of a Late Cretaceous specimen, so we were confident in assuming that they were fossil *Podocarpus*.

The limited morphological differentiation among *Podocarpus* species makes interpreting the placement of our fossil specimens difficult. However, the spatio-temporal pattern that we observe is consistent with the biogeographic history of *P. Podocarpus* [35]. *In our tip-calibrated tree, the 67*.*2 Ma Colombian specimen was placed as sister to the entire subgenus P. Podocarpus*, suggesting a lineage now extinct in the Neotropics. The Miocene Peruvian specimens were placed with the early-diverging Chilean species, *P. salignus*. The four Panamanian specimens were the youngest fossils in our analysis. As would be predicted, three specimens (3.55, 4.25 and 10.15 Ma) were placed with high support with the main Neotropical lineage (*P. sellowii, P. coriaceus, P. brasiliensis, P. urbanii, P. oleifolius*, and *P. matudae*).

However, the youngest Panamanian specimen (2.05 Ma) was highly corroded, with few visible features on the sacci or corpus. Its unlikely placement with *P. salignus* is likely the result of this poor preservation. Its placement, and the placement of the four Miocene (12 Ma) Peruvian specimens, with the Chilean species *P. salignus* had poor support. However, excluding the specimen did not affect the placement of the Peruvian fossils (Supplemental Fig. S4). The low node support may instead reflect the uncertainty in the placement of *P. salignus* within *P. Podocarpus* [24, 35].

## 5 Conclusions

Palynologists have long used the morphological conservation of morphological features within clades to classify pollen taxa. However, our results suggest that an even deeper evolutionary history is contained within pollen morphology. The vocabulary of palynology is vast [32], but it does not capture all the gradations and permutations possible in pollen wall structure and ornamentation. Deep learning connects morphology to molecular phylogenies, learning evolutionary distances from existing phylogenies and transforming morphological classification features. Features are transformed from characters that define individual clades to the synapomorphies that link them.

Deep learning is adaptive to the constraints of paleontological data, including incomplete or unresolved phylogenies and fossil morphotypes derived from extinct taxa. Novelty detection recognizes new morphologies and expands our knowledge of a clade’s morphospace. We can explicitly quantify our uncertainty. The abundance of fossil pollen in geological samples also means that phylogenetic determinations can be made with populations of specimens, not just individual specimens. Applied at a broad scale, phylogenetically-informed machine learning models will allow palynologists to harness the evolutionary information within the fossil record and identify taxon origins and extinctions that would otherwise go unrecognized.

## Acknowledgments

We thank Carlos Jaramillo for sharing fossil and reference pollen samples, Timme Donders for providing additional reference pollen samples, Andrew Leslie for providing the *Podocarpus* phylogeny, and Milton Tan for feedback on the manuscript. We thank Ingrid Romero and Michael Urban for imaging the majority of the material used and cited in this analysis. Support for MEA and SWP was provided by a 2022 National Center for Supercomputing Applications Faculty Fellowship to SWP and the University of Illinois Tom L. Phillips Fund for Paleobotany.

## Data availability

The Airyscan images used in this study were submitted to the Illinois Databank (https://doi.org/10.13012/B2IDB-8817604_V1). The codebase is publicly available on GitHub (DOI: 10.5281/zenodo.8128443) and can be accessed using this link: https://github.com/paleopollen/Novel_Pollen_Phylogenetic_Placement.

