## Supplementary Information for "Deep Learning Approaches to the Phylogenetic Placement of Extinct Pollen Morphotypes"

Surangi W. Punyasena

### **RESULTS**

**Baseline classification accuracy**

**Fig. S1.** Following are the confusion matrices of the classification accuracies for the modern *Podocarpus* specimens for five training/validation splits and their average. Shown are H-CNN (upper left), C-CNN (upper right), P-CNN (lower left) and FM (lower right). Rows represent the true taxon while columns represent the model’s predictions.

Split 1

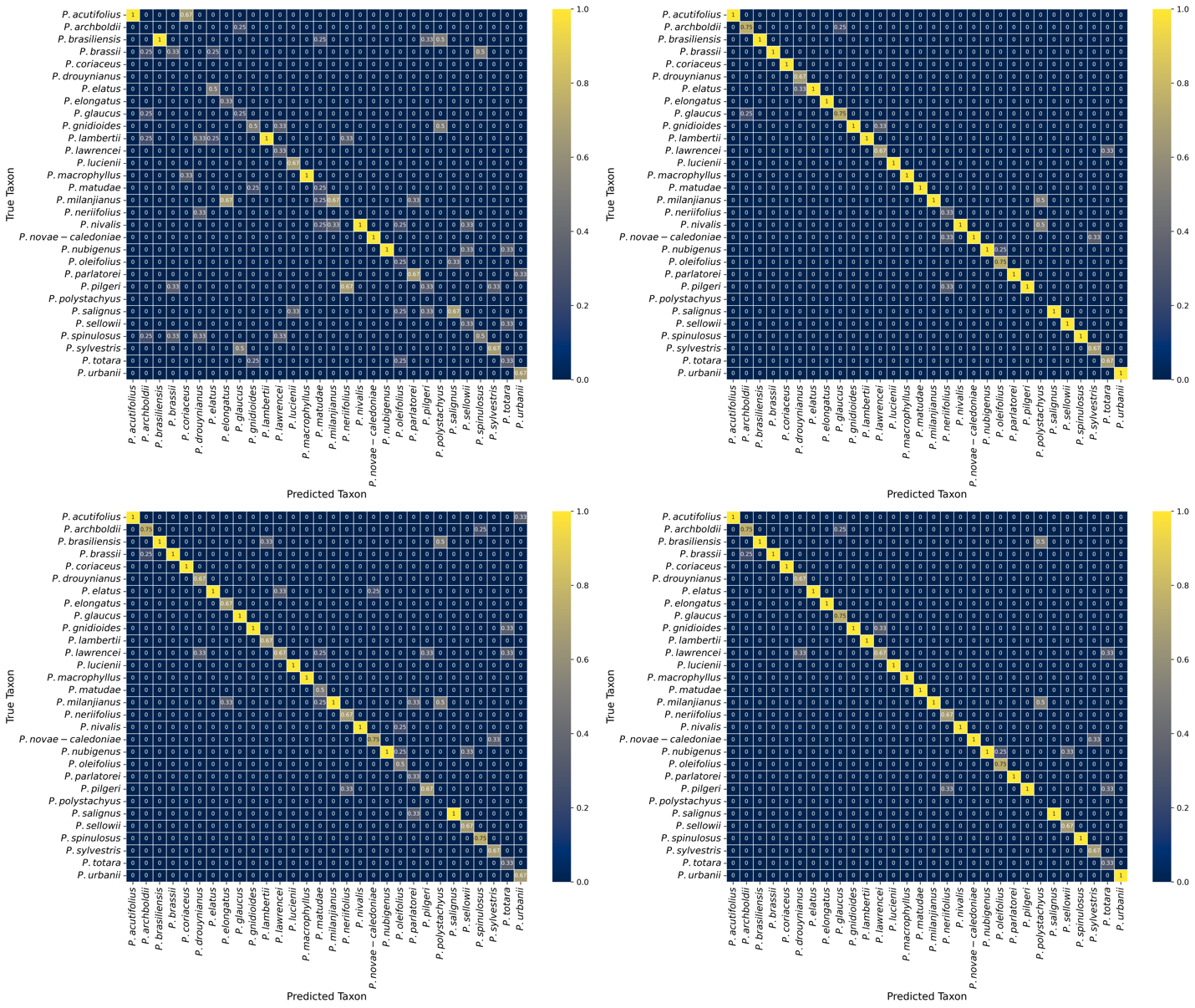

Split 2

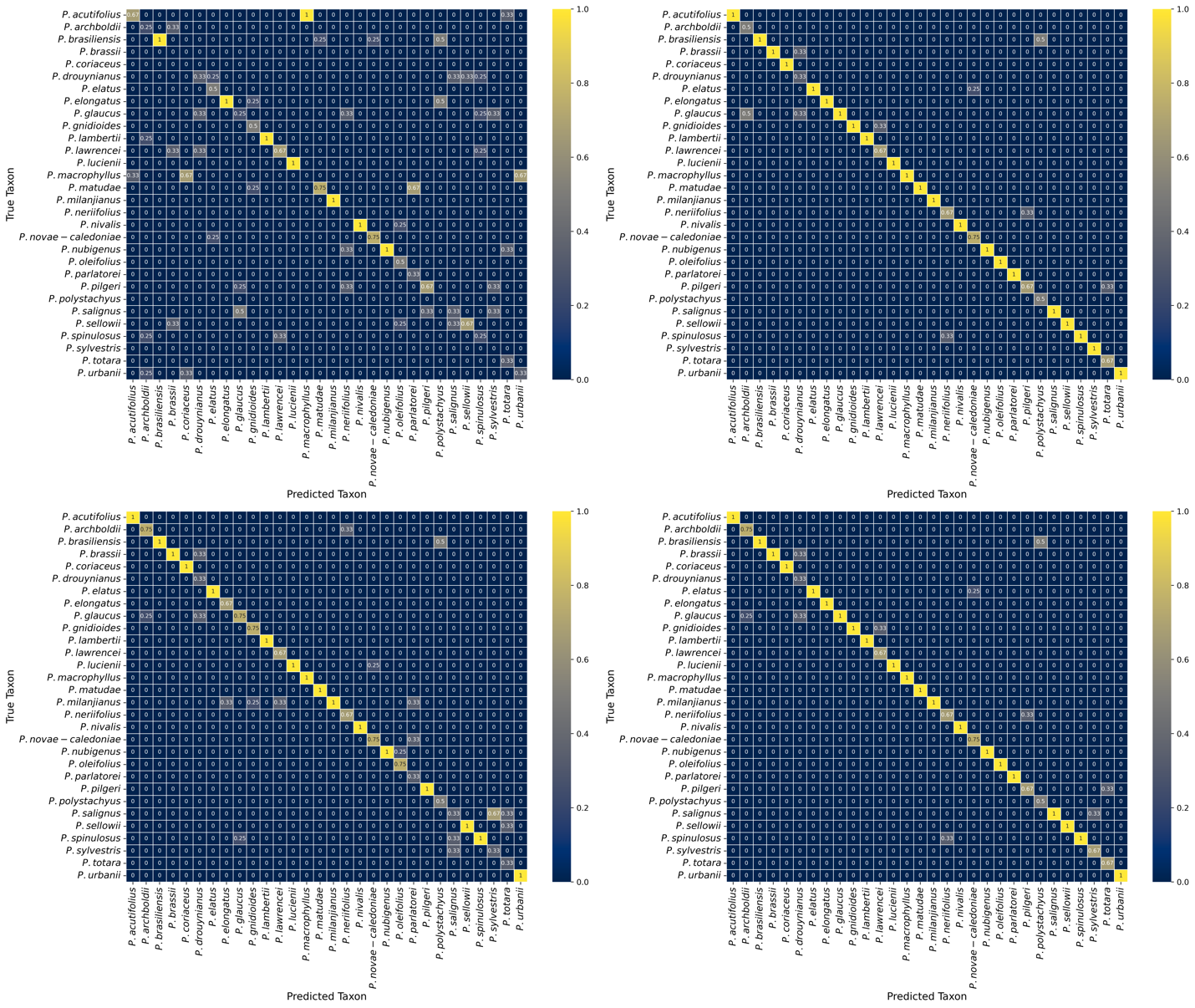

Split 3

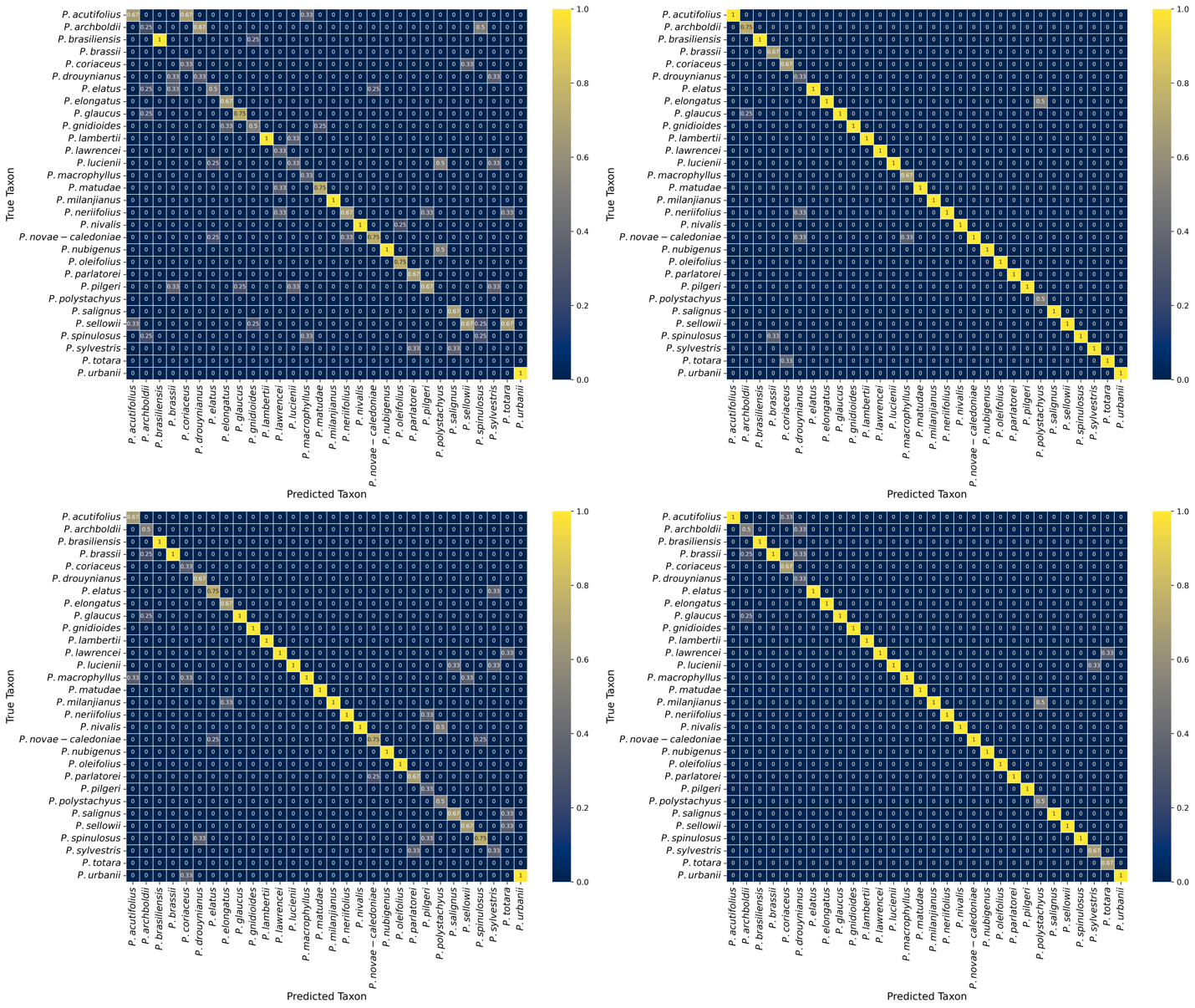

Split 4

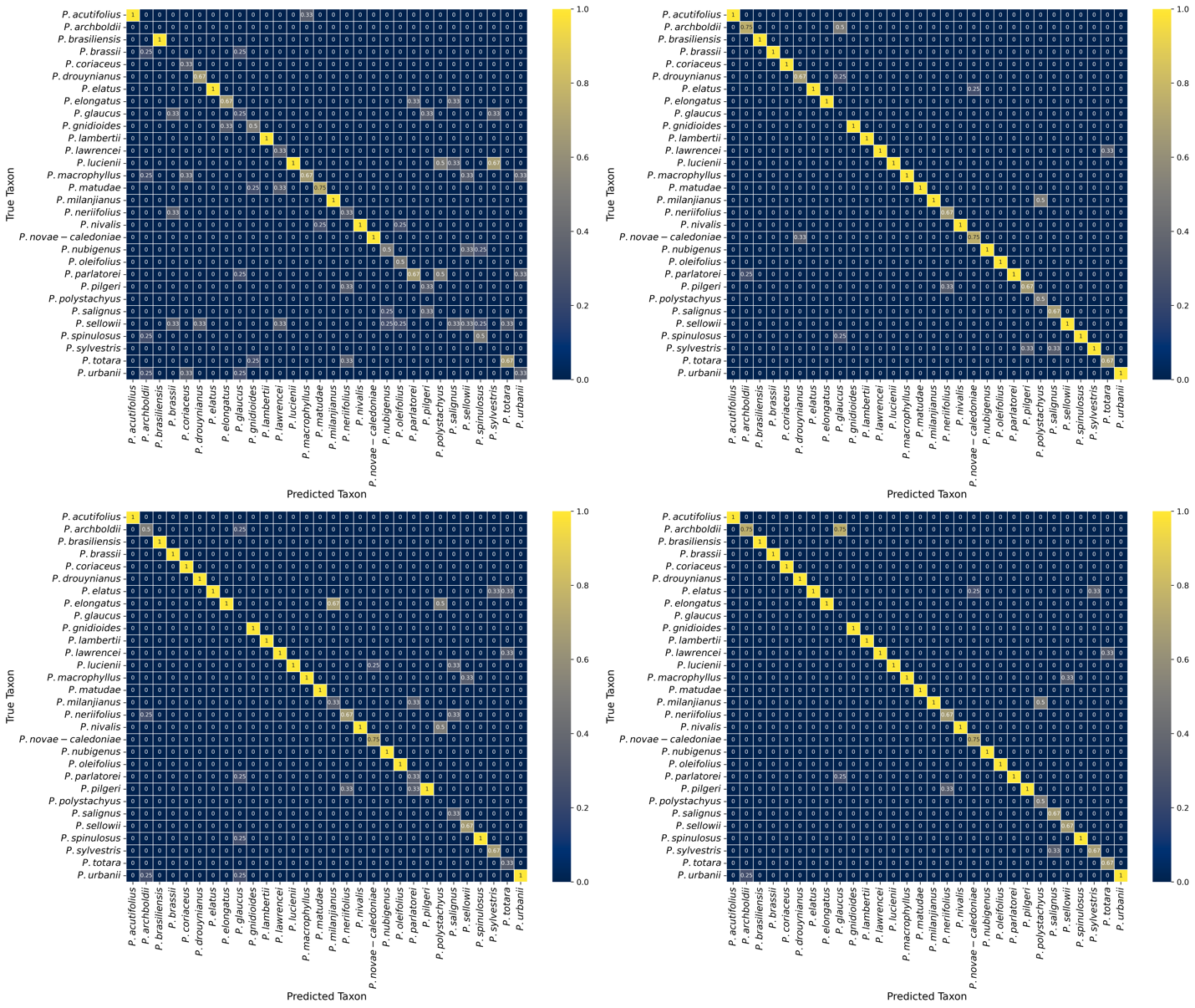

Split 5

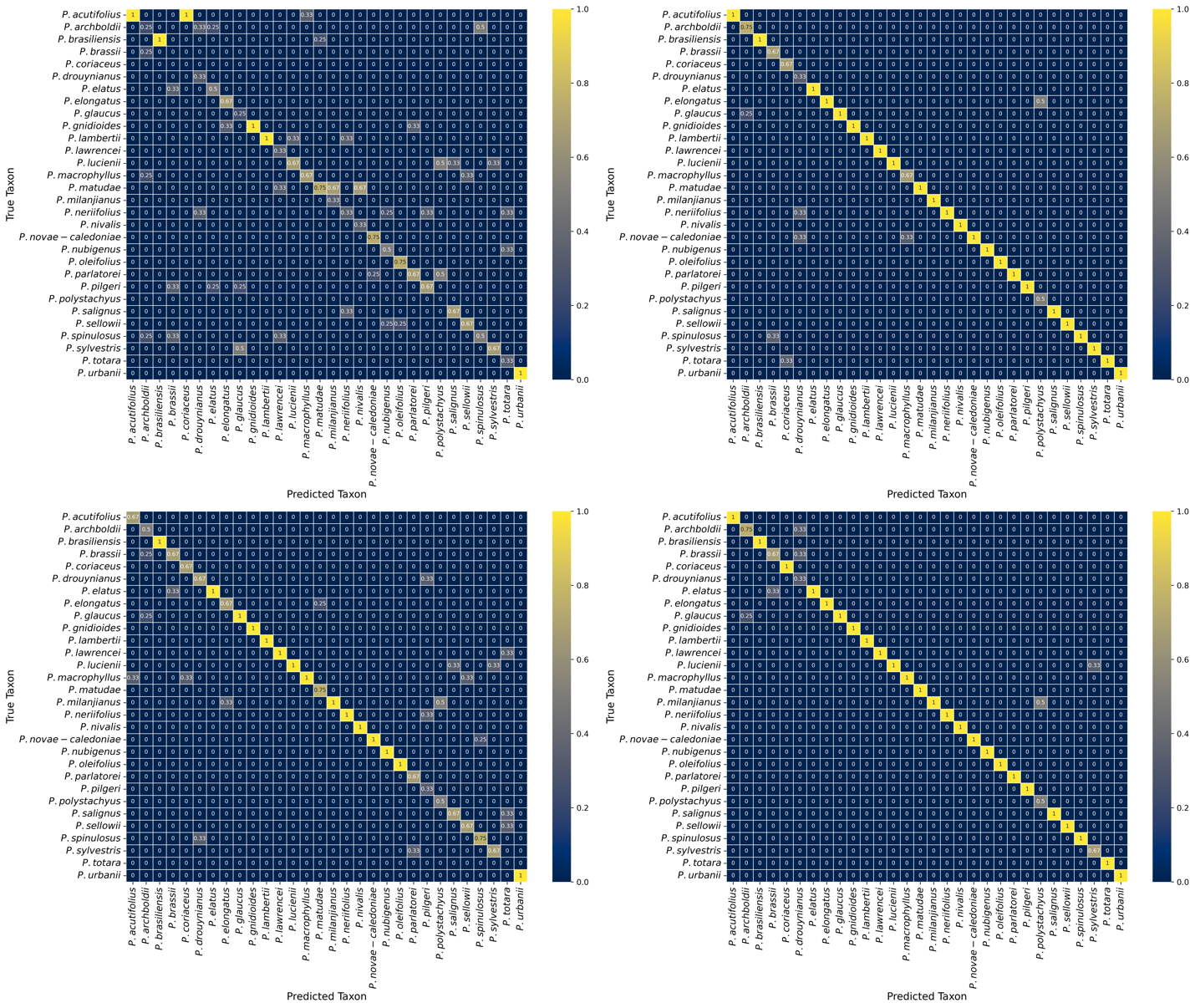

Average of all five splits

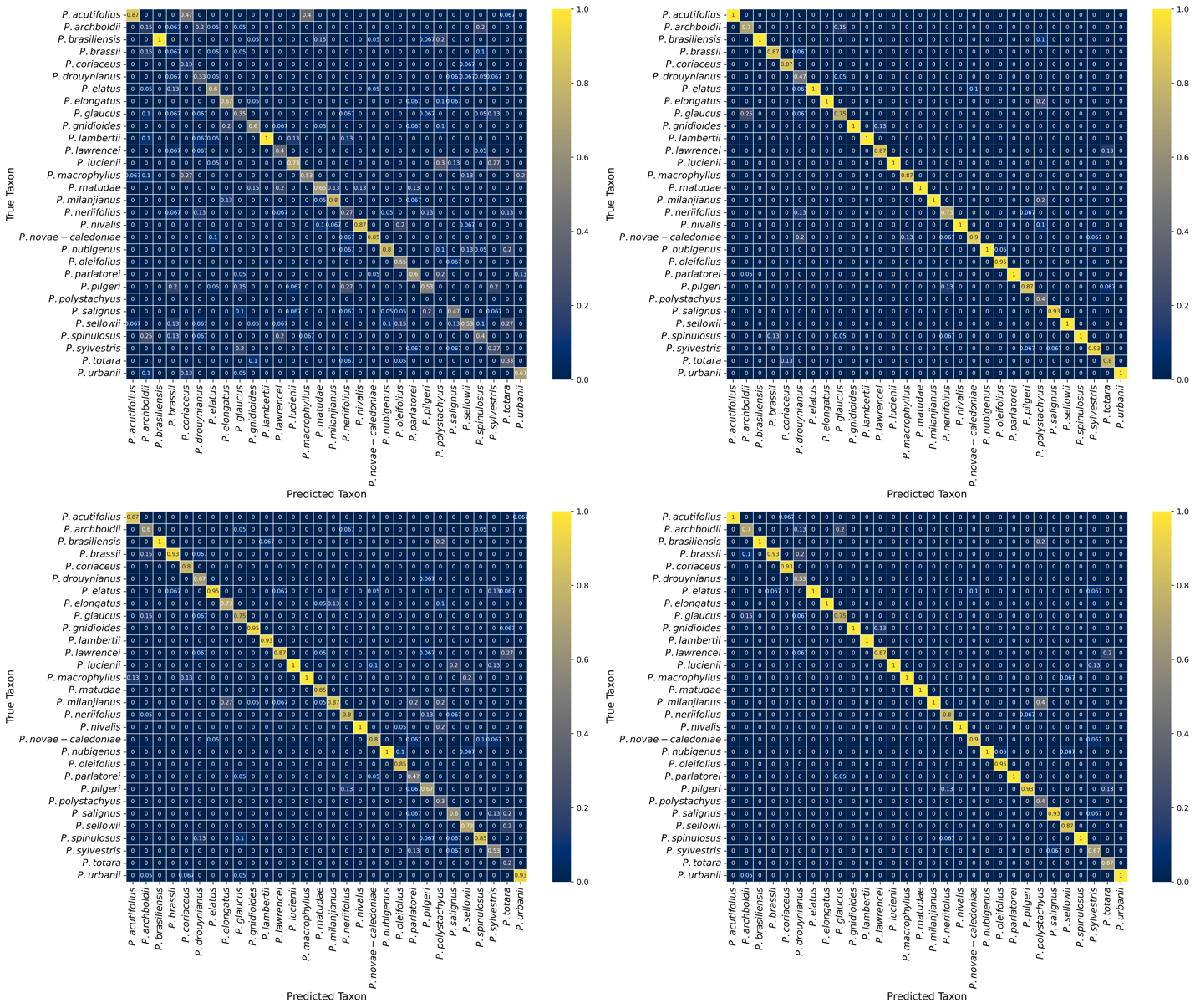

**Novel taxon detection**

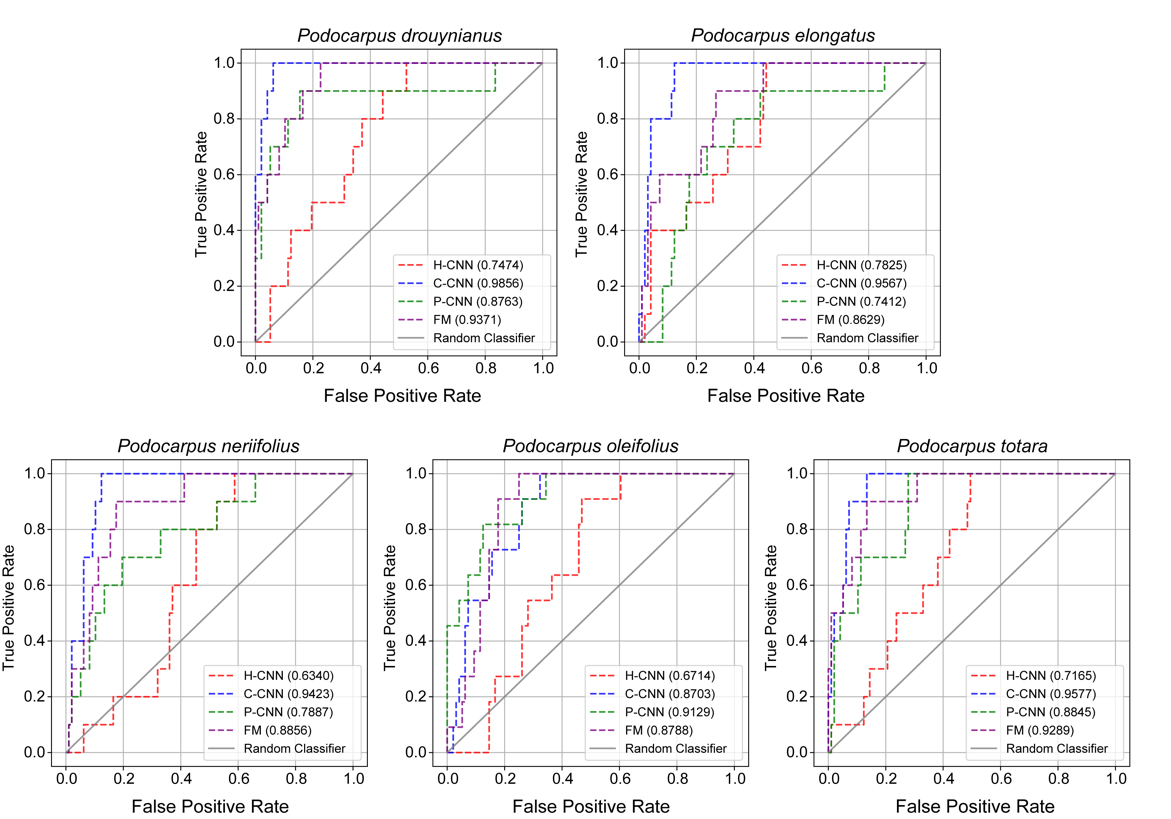

**Fig. S2.** Receiver operating characteristic (ROC) curves plotted for each of the five novelty detection experiments, and their corresponding areas ranging from 0 to 1. The curves were computed using the four models, namely the holistic CNN (H-CNN), cross-sectional CNN (C-CNN), patch CNN (P-CNN), and fused model (FM). The greater the area under the curve, the better the model discriminates among known and novel pollen types.

**Phylogenetic placement of pseudo-novel specimens**

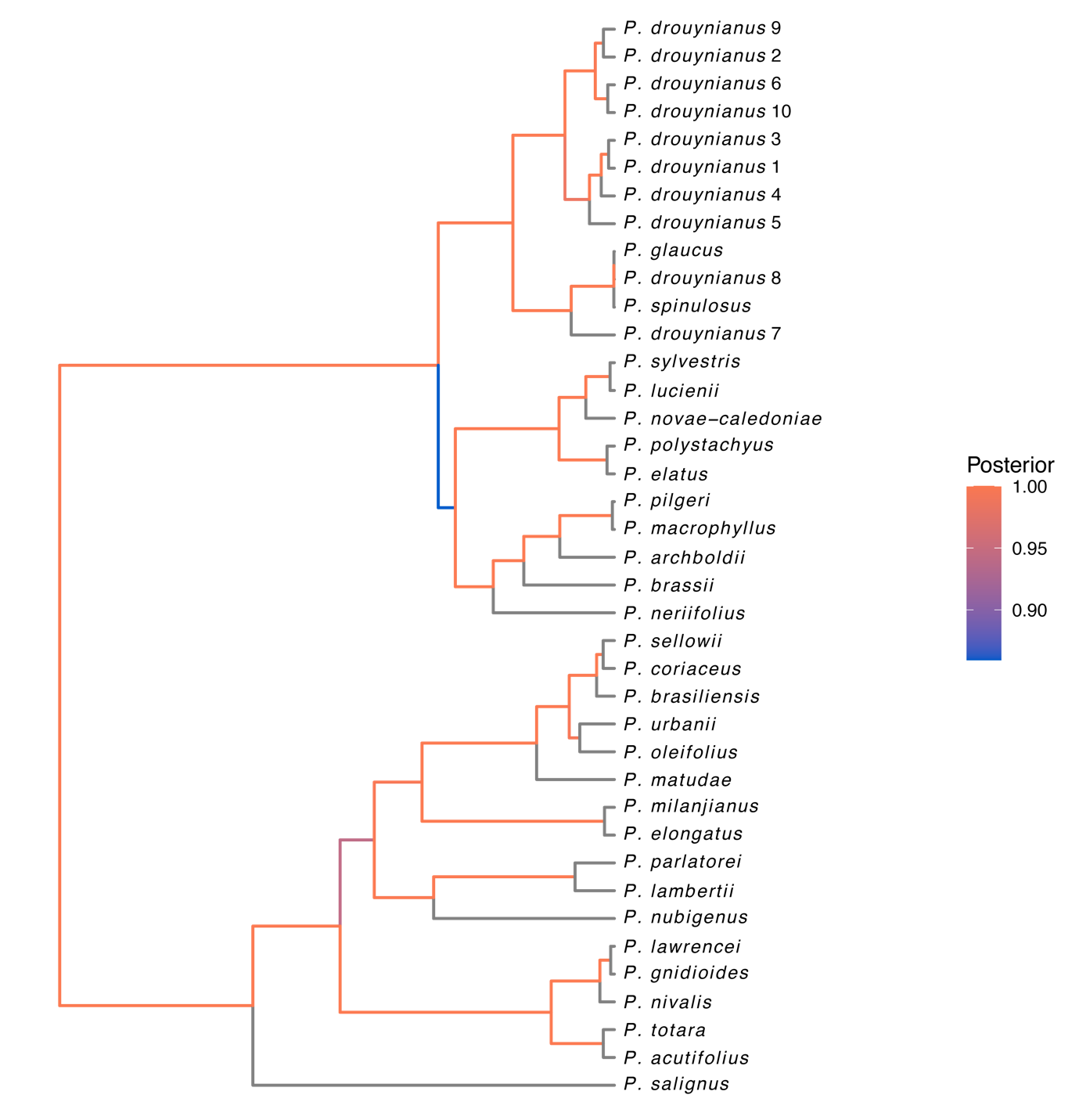

**Fig. S3.1.** Bayesian tree for *Podocarpus drouynianus.* The tree replicates the topology in Leslie et al. (2018). Branches are coloured according to posterior probability. All ten pseudo-novel specimens are accurately placed with *P. glaucus* and *P. spinulosus*, with high support (P=1) for this clade.

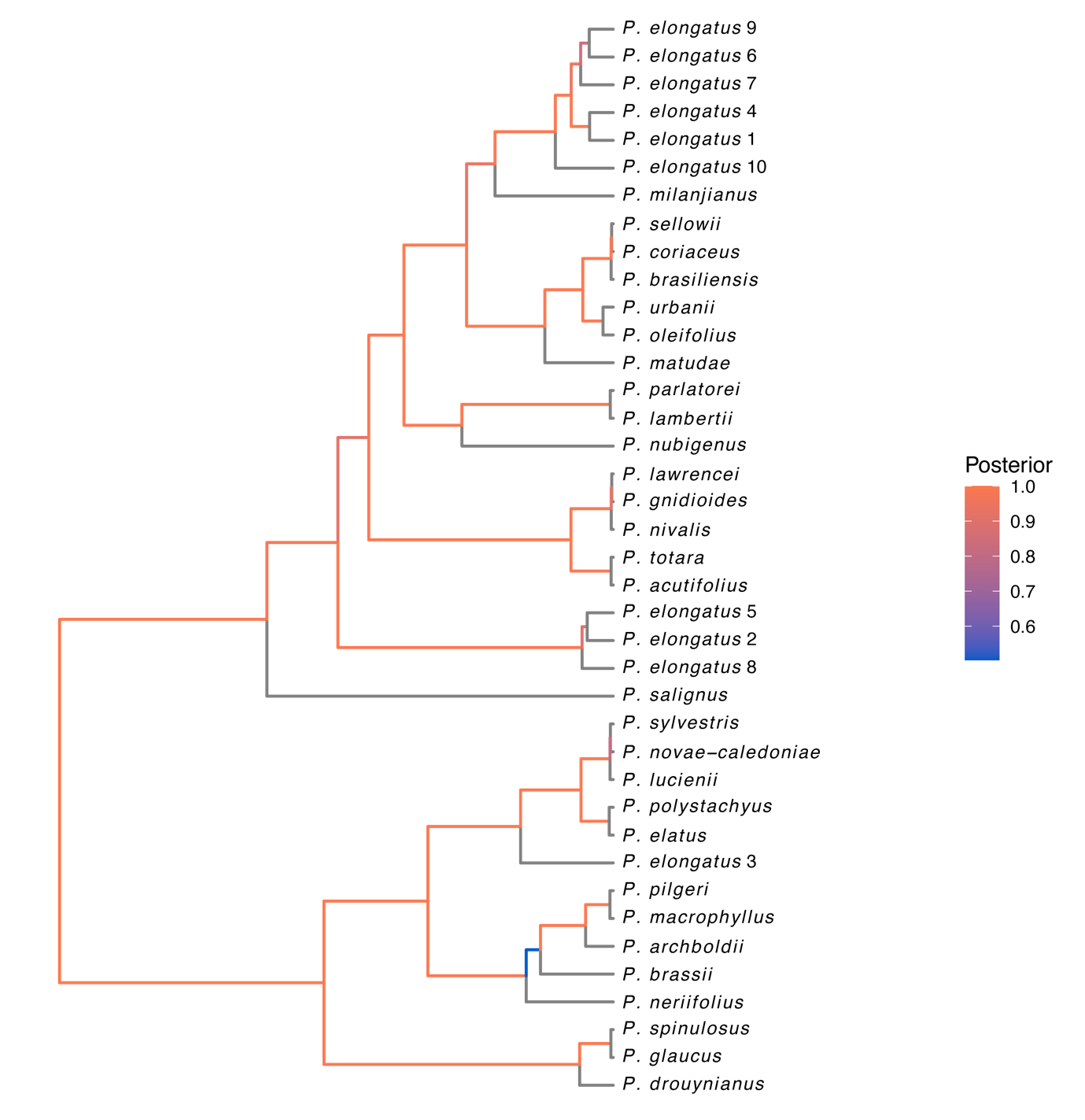

**Fig. S3.2.** Bayesian tree for *Podocarpus elongatus.* The tree replicates the topology in Leslie et al. (2018). Branches are coloured according to posterior probability. Of the ten pseudo-novel specimens, six are accurately placed with *P*. *milanjianus*, with relatively high support (P=0.935) for this clade. Three other specimens are embedded elsewhere in *Podocarpus* subgenus *Podocarpus*, and the remaining one is placed within subgenus *Foliolatus*.

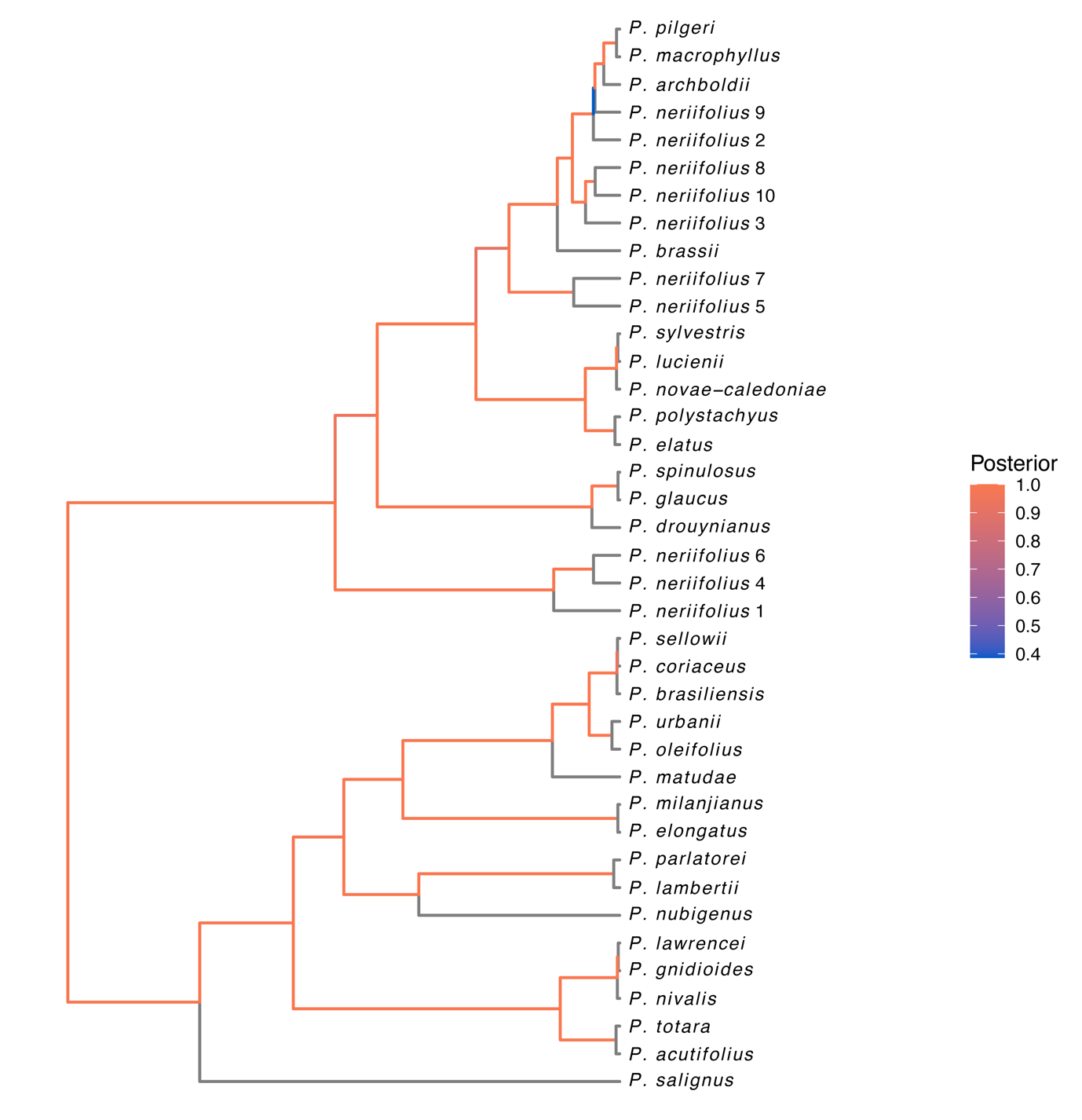

**Fig. S3.3.** Bayesian tree for *Podocarpus neriifolius.* The tree replicates the topology in Leslie et al. (2018). Branches are coloured according to posterior probability. Of the ten pseudo-novel specimens, seven are accurately embedded in the subclade comprising *P. brassii*, *P. archboldii*, *P. macrophyllus*, and *P. pilgeri*, with relatively high support (P=0.910) for this clade. The remaining three specimens were placed as sister to subgenus *Foliolatus*.

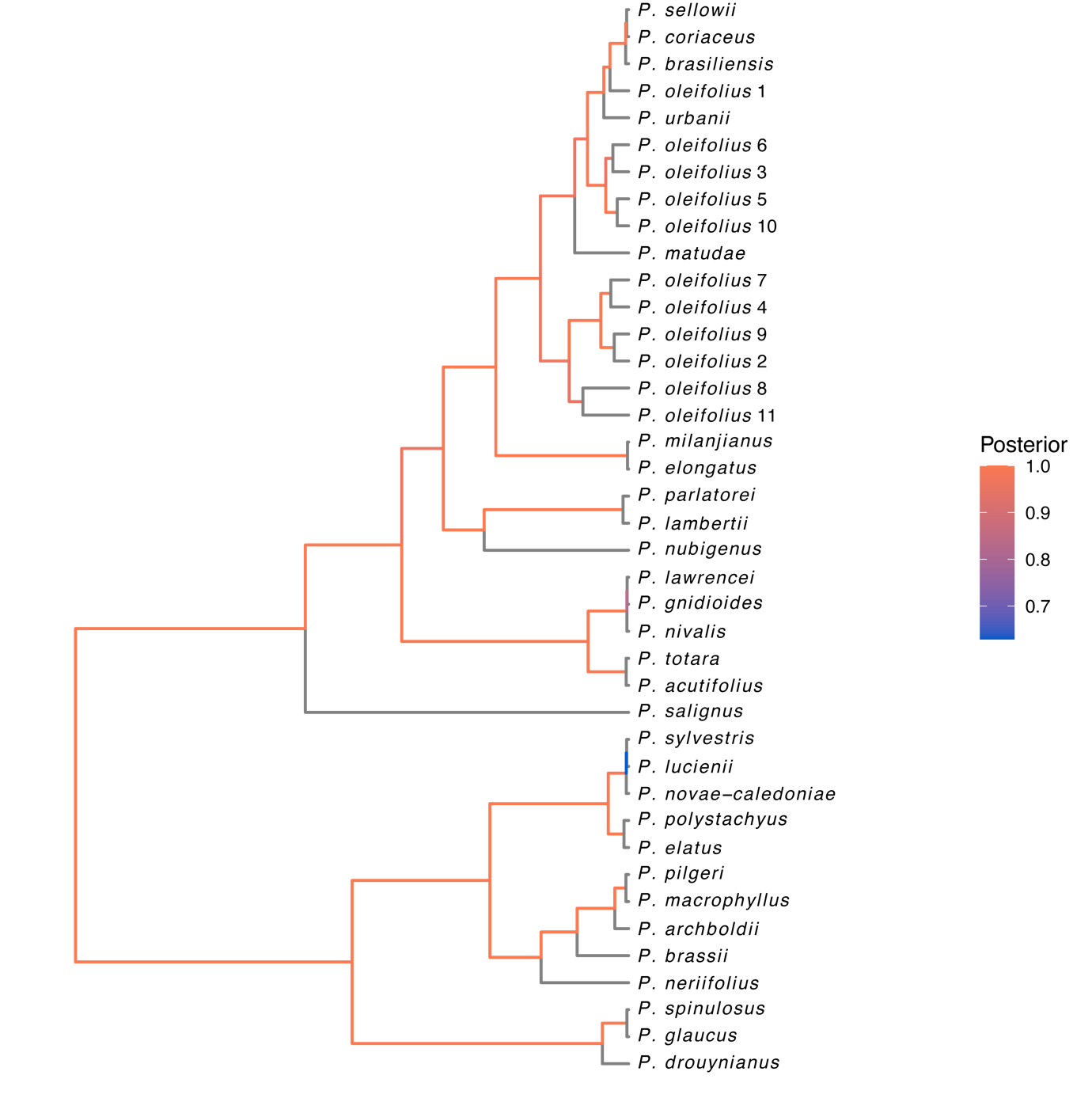

**Fig. S3.4.** Bayesian tree for *Podocarpus oleifolius.* The tree replicates the topology in Leslie et al. (2018). Branches are coloured according to posterior probability. All 11 pseudo-novel specimens are accurately placed with the Neotropical clade comprising *P. coriaceus*, *P. brasiliensis*, *P. sellowii*, *P. urbanii*, and *P. matudae*, with high support (P=1) for this clade.

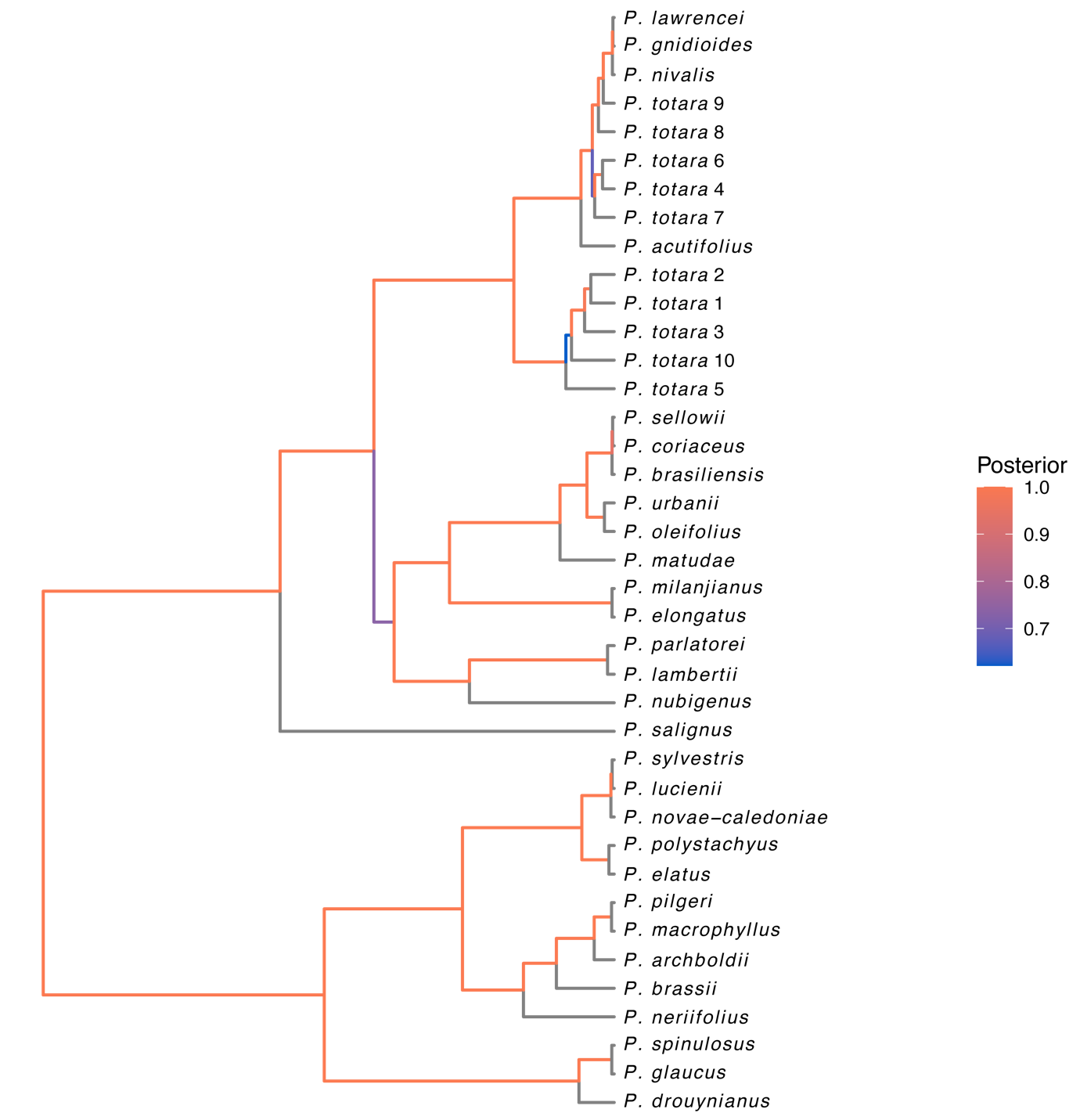

**Fig. S3.5.** Bayesian tree for *Podocarpus totara.* The tree replicates the topology in Leslie et al. (2018). Branches are coloured according to posterior probability. All ten pseudo-novel specimens are accurately placed with the Australasian clade comprising *P. acutifolius*, *P. gnidioides*, *P. lawrencei*, and *P. nivalis,* with high support (P=1) for this clade.

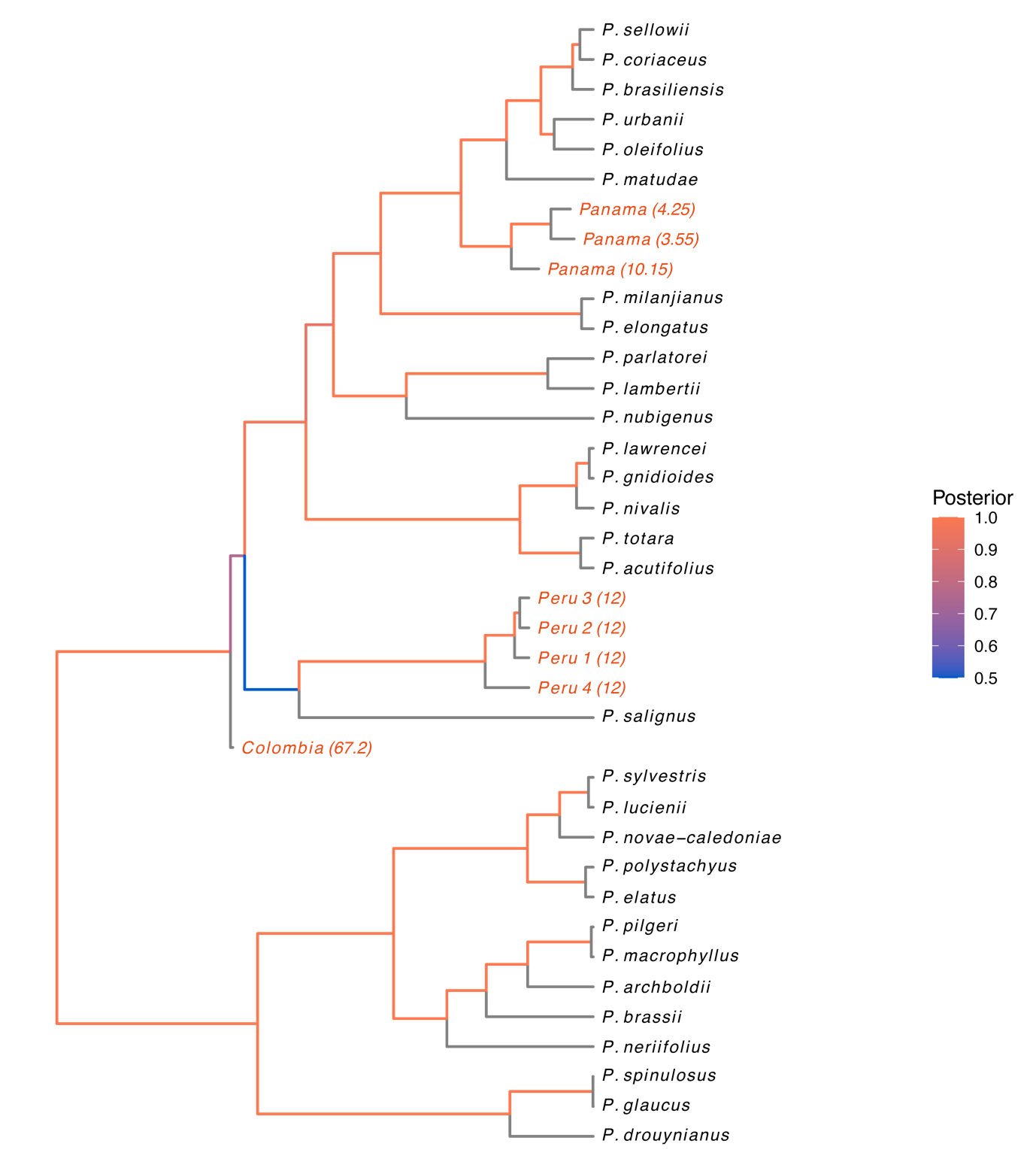

**Fig. S4.** Combined tip-calibrated fossil tree, excluding the youngest Panamanian (2.05 Ma) specimen, simulated under the fossilized birth-death process (FBDP) using the MLP-transformed features. The tree topology replicates that of the reference phylogeny proposed by Leslie et al. (2018). Branch lengths were estimated based on our fossil tip ages. The three Panamanian specimens are placed as sister to the Central-South American tropical clade formed by *P. sellowii*, *P. coriaceus*, *P. brasiliensis*, *P. urbanii*, *P. oleifolius*, and *P. matudae*. The four Peruvian specimens are placed with the Chilean *P. salignus*. Finally, the late Cretaceous Colombian specimen is placed as sister to the subgenus P. Podocarpus.

**MLP architecture**

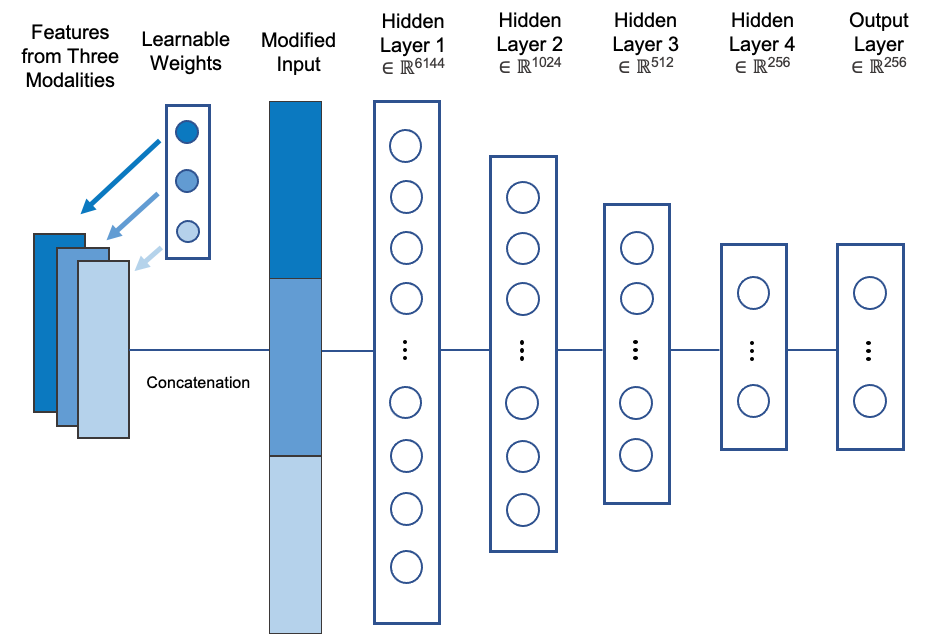

**Fig. S5.** Overview of the MLP architecture. Dropout layer (with dropout rate = 0.2) was inserted between each two consecutive layers to avoid overfitting.

**Baseline classification accuracy**

**Table S1.** Five-fold cross-validation classification accuracy for the holistic CNN (H-CNN), cross-sectional CNN (C-CNN), patch CNN (P-CNN), and fused model (FM) of the *Podocarpus* dataset, with their corresponding mean and standard deviation.

| Split Number | Classification Accuracy (%) | | | |
| --- | --- | --- | --- | --- |
|  | **H-CNN** | **C-CNN** | **P-CNN** | **FM** |
| 1 | 52.00 | 89.00 | 78.00 | 88.00 |
| 2 | 52.00 | 90.00 | 81.00 | 90.00 |
| 3 | 58.00 | 93.00 | 79.00 | 92.00 |
| 4 | 56.00 | 88.00 | 80.00 | 88.00 |
| 5 | 57.00 | 93.00 | 81.00 | 94.00 |
| **Mean** | **55.00** | **90.60** | **79.80** | **90.40** |
| **s.d.** | **2.83** | **2.30** | **1.30** | **2.61** |

**Novel taxon detection**

**Table S2.** AUROC values (ranging between 0 and 1) for the pseudo-novel *Podocarpus* species under the four classification models. Values are also reported in Supplemental Fig. S2.

| **Species** | **C-CNN** | **H-CNN** | **P-CNN** | **FM** |
| --- | --- | --- | --- | --- |
| *Podocarpus drouynianus* | 0.9856 | 0.7474 | 0.8763 | 0.9371 |
| *Podocarpus elongatus* | 0.9567 | 0.7825 | 0.7412 | 0.8629 |
| *Podocarpus neriifolius* | 0.9423 | 0.6340 | 0.7887 | 0.8856 |
| *Podocarpus oleifolius* | 0.8703 | 0.6714 | 0.9129 | 0.8788 |
| *Podocarpus totara* | 0.9577 | 0.7165 | 0.8845 | 0.9289 |
| **Mean** | **0.9425** | **0.7104** | **0.8407** | **0.8987** |
| **s.d.** | **0.0433** | **0.0591** | **0.0725** | **0.0325** |

#### MLP Learned Weights

**Table S3.** Learned weights for MLP concatenation of CNN features. Five runs per species per CNN model are listed.

| **Pseudo-novel taxon** | **H-CNN weights** | **C-CNN weights** | **P-CNN weights** |
| --- | --- | --- | --- |
| *Podocarpus drouynianus* | 0.3320  0.3317  0.3323  0.3316  0.3313 | 0.3353  0.3361  0.3356  0.3361  0.3363 | 0.3327  0.3322  0.3320  0.3323  0.3324 |
| *Podocarpus elongatus* | 0.3313  0.3318  0.3320  0.3322  0.3327 | 0.3365  0.3361  0.3357  0.3356  0.3348 | 0.3321  0.3321  0.3323  0.3323  0.3324 |
| *Podocarpus neriifolius* | 0.3318  0.3327  0.3319  0.3320  0.3319 | 0.3361  0.3349  0.3358  0.3358  0.3360 | 0.3321  0.3324  0.3323  0.3321  0.3320 |
| *Podocarpus oleifolius* | 0.3315  0.3317  0.3318  0.3318  0.3316 | 0.3360  0.3362  0.3358  0.3360  0.3363 | 0.3325  0.3321  0.3324  0.3322  0.3321 |
| *Podocarpus totara* | 0.3312  0.3314  0.3309  0.3315  0.3311 | 0.3395  0.3384  0.3388  0.3389  0.3390 | 0.3293  0.3302  0.3303  0.3296  0.3299 |

### **MATERIALS AND METHODS**

#### Modern specimen information

**Table S4.** Modern *Podocarpus* reference specimens. Images from Punyasena et al. (2023).

| **Plate Numbers** | **Species** | **Number of Specimens** | **Geographic Range** | **Collection** | **Slide ID** |
| --- | --- | --- | --- | --- | --- |
| 1-2 | *Podocarpus acutifolius* | 10 | Aust | STRI - Graham | 1675959 |
| 3-4 | *Podocarpus archboldii* | 11 | Aust | STRI - Graham | 19848 |
| 5-6 | *Podocarpus brasiliensis* | 14 | Neo | STRI - CTPA | 896 |
| 7-8 | *Podocarpus brassii* | 10 | Aust | STRI - Graham | NA |
| 9-10 | *Podocarpus coriaceus* | 10 | Neo | STRI - Graham | 845414 |
| 11-12 | *Podocarpus drouynianus* | 10 | Aust | STRI - Graham | 19836 |
| 13-14 | *Podocarpus elatus* | 11 | Aust | STRI - Graham | 19837 |
| 15-16 | *Podocarpus elongatus* | 10 | Afr | Utrecht | 1540 |
| 17-18 | *Podocarpus glaucus* | 11 | Aust/Indomalay | STRI - Graham | 19847 |
| 19-20 | *Podocarpus gnidioides* | 12 | Aust | STRI - Graham | 22388 |
| 21-22 | *Podocarpus lambertii* | 10 | Neo | STRI-CTPA | 898 |
| 23-24 | *Podocarpus lawrencei* | 10 | Aust | STRI - Graham | 19839 |
| 25-26 | *Podocarpus lucienii* | 10 | Aust | STRI - Graham | 22391 |
| 27-28 | *Podocarpus macrophyllus* | 10 | Palearc | STRI - Graham | 63458 |
| 29-30 | *Podocarpus matudae* | 11 | Neo | STRI - Graham | 10584 |
| 31-32 | *Podocarpus milanjianus* | 10 | Afr | Utrecht | NA |
| 33-34 | *Podocarpus neriifolius* | 10 | Aust/Indomalay | STRI - Graham | 19844 |
| 35-36 | *Podocarpus nivalis* | 10 | Aust | STRI - Graham | 12961 |
| 37-38 | *Podocarpus novae-caledoniae* | 11 | Aust | STRI - Graham | 22392 |
| 39-40 | *Podocarpus nubigenus* | 11 | Neo | STRI - CTPA | 1392 |
| 41-42 | *Podocarpus oleifolius* | 11 | Neo | STRI - CTPA | 1393 |
| 43-44 | *Podocarpus parlatorei* | 9 | Neo | STRI - Graham | 8345 8349 |
| 45-46 | *Podocarpus pilgeri* | 10 | Indomalay | STRI - Graham | 19845 |
| 47-48 | *Podocarpus polystachyus* | 5 | Aust/Indomalay | Utrecht | 19849 |
| 49-50 | *Podocarpus salignus* | 10 | Neo | STRI - Graham | 8346 |
| 51-52 | *Podocarpus sellowii* | 10 | Neo | STRI - CTPA | 1395 |
| 53-54 | *Podocarpus spinulosus* | 12 | Aust | STRI - Graham | 19841 |
| 55-56 | *Podocarpus sylvestris* | 10 | Aust | STRI - Graham | 19833 |
| 57-58 | *Podocarpus totara* | 10 | Aust | STRI - Graham | 19843 |
| 59-60 | *Podocarpus urbanii* | 10 | Neo | STRI - Graham | 427685 |

STRI = Smithsonian Tropical Research Institute

**Table S5.** *Podocarpidites* fossil specimen slide and sample details. Ages were calculated following the maximum likelihood-based biostratigraphic method described in Punyasena et al. (2012). The specimens include pollen types previously described in Jaramillo et al. (2014), Martinez et al. (2020), and Carvalho et al. (2020). Images from Punyasena et al. (2023).

| **Plate Numbers** | **Specimen** | **Slide number and Information** | **Country** | **Latitude** | **Longitude** | **Min**  **Age**  **(Mya)** | **Max**  **Age**  **(Mya)** |
| --- | --- | --- | --- | --- | --- | --- | --- |
| F1-2 | Colombia (67.2) | PG1-180m | Colombia | 6.55 | -73.7 | 67.1 | 67.2 |
| F3-4 | Peru 1 (12) | ID 36583 San Miguel Field Station, Peru, member B | Peru | -14.6663 | -71.2832 | 10 | 12 |
| F5-6 | Peru 2 (12) | ID 36584 Acocunca North, Member B | Peru | -14.7276 | -71.2674 | 10 | 12 |
| F7-8 | Peru 3 (12) | ID 39456 El Descanso, Member B | Peru | -14.6624 | -71.2758 | 10 | 12 |
| F9-10 | Peru 4 (12) | ID 39460 El Descanso, Member B | Peru | -14.6624 | -71.2758 | 10 | 12 |
| F11-12 | Panama (10.15) | PPP 900 Fm. Tuira, Rio Tuira, Darien, Panama | Panama | 8.13188889 | -77.658 | 10.15 | 10.15 |
| F13-14 | Panama (4.25) | CJ-87-27-01 Fm. Cayo Agua, Bocas del Toro, Panama | Panama | 9.17641667 | -82.058 | 4.25 | 4.25 |
| F15-16 | Panama (3.55) | PPP 02182 Fm. Escudo Veraguas,  Bocas del Toro, Panama | Panama | 9.087969 | -81.53611 | 3.55 | 3.55 |
| F17-18 | Panama (2.05) | CJ-87-11-01 Fm. Escudo Veraguas, Bocas del Toro, Panama | Panama | 9.10327778 | -81.57466 | 2.05 | 2.05 |

**Table S6.** Table detailing the MLP architecture. A rectified linear activation unit (ReLU) was inserted between each two consecutive layers.

| **Layer** | **Input dim** | **Output dim** |
| --- | --- | --- |
| Attentional Weighting | 6144 | 6144 |
| Fully Connected (fc1) | 6144 | 1024 |
| Fully Connected (fc2) | 1024 | 512 |
| Dropout 1 (P = 0.2) | 512 | 512 |
| Fully Connected (fc3) | 512 | 256 |
| Dropout 2 (P = 0.2) | 256 | 256 |
| Fully Connected (fc4) | 256 | 256 |
